# Large haplotypes linked to climate and life history variation in divergent lineages of Atlantic salmon (*Salmo salar*)

**DOI:** 10.1101/2025.05.23.655798

**Authors:** S.J. Lehnert, T. Kess, P. Bentzen, N. Barson, S. Lien, J.B. Dempson, I.R. Bradbury

## Abstract

Advances in sequencing are revealing that linked genomic architectures, enabling the evolution of co-adapted alleles at multiple loci, often shape complex phenotypes. Several recent studies have identified such architectures (e.g., chromosomal rearrangements and supergenes) contributing to adaptation or divergence across diverse species, from plants to mammals. Specifically, within Atlantic salmon (*Salmo salar*), genomic studies are revealing large haplotypes and structural variants that may underpin local adaptation in the species. Using data from >4000 individuals from 134 locations spanning the North Atlantic Ocean, we identify a large (∼3-Mbp) genomic region on Ssa18 showing patterns of differentiation and linkage disequilibrium indicative of a large haplotype block containing three divergent haplotypes (herein A, B, and C haplotypes). In Europe, haplotypes A and B were common, whereas A and C were more common within North America, suggesting a shared ‘ancestral’ A haplotype, with different continent-specific alternative haplotypes. Data support independent origins of divergent haplotypes in each continent, as well as signals of trans-oceanic introgression of haplotypes. Haplotype frequency is strongly associated with latitude, climate, and life history (smolt age); however, the strength and direction of these relationships vary across continents. Overall analyses were consistent with other studies that identify chromosomal rearrangements; however, long read sequence data could not confirm evidence of a structural variant, although an ancestral fusion polymorphism may explain the formation and maintenance of the observed haplotypes. Our study contributes to ongoing efforts to understand the evolutionary role of linked genomic architecture in Atlantic salmon and its significance in salmonid diversification.

## Introduction

Understanding the underlying genetic variation that contributes to adaptation remains a central goal in evolutionary biology. While the last decade of genomic work has focused on variation in single nucleotide polymorphisms (SNPs), advances in sequencing technology are revealing other sources of genomic variation that shape ecological and evolutionary processes (Huang et al., 2020; Mérot et al., 2020; Ishigohoka et al., 2024). Recent genomic studies have highlighted how linked genomic architectures, which facilitate the evolution of co-adapted alleles at multiple loci, contribute to adaptation across taxa (Schwander et al., 2014; Wellenreuther & Bernatchez, 2018; Mérot et al., 2020). This phenomenon has been observed in ecotype formation across a wide range of species, from plants to mammals (Huang et al., 2020; Hager et al., 2022). Selection on these tightly linked loci can lead to suppressed recombination that enable multiple loci to be inherited as a single haplotype, thus enabling simple Mendelian inheritance of complex phenotypes that can be maintained in the face of gene flow (Charlesworth & Charlesworth, 1973; Mérot et al., 2020; Oomen et al., 2020). Different mechanisms can lead to the formation of such haplotypes (Schwander et al., 2014; Thompson & Jiggins, 2014; Mérot et al., 2020; Ishigohoka et al., 2024), often (but not always) involving large scale chromosomal changes such as inversions, fusions, and translocations (Charlesworth & Charlesworth, 1973; Wellenreuther & Bernatchez, 2018; Lehnert et al., 2019; Huang et al., 2020; Harringmeyer & Hoekstra, 2022; Stenløkk et al., 2022).

Within salmonids, several recent studies have identified linked architectures contributing to adaptation or divergence, such as the identification of large Y-chromosome haplotypes driving age of maturity in male Chinook salmon (*Oncorhynchus tshawytscha*) (McKinney et al., 2020; McKinney et al., 2021), haplotypes in regions of low recombination associated with divergent lineages of Arctic charr (*Salvelinus alpinus*) (Dallaire et al., 2024), and chromosomal rearrangements associated with environment in Atlantic salmon (*Salmo salar*) (Stenløkk et al., 2022; Watson et al., 2022). These regions can range in size from smaller scale haplotype blocks such as a ∼0.14-Mb region influencing migration timing in Chinook salmon (Thompson et al., 2020) to large structural changes such as a 55-Mb double-inversion supergene controlling life history in rainbow trout (*O. mykiss*) (Pearse et al., 2019).

These examples highlight the role of tightly linked loci in driving extensive diversity observed within salmonids. Like many other salmonids, Atlantic salmon exhibits an anadromous life history and displays strong natal philopatry, resulting in high levels of genetic structure across multiple spatial scales (Moore et al., 2014; Bradbury et al., 2018; Lehnert et al., 2020). Given that Atlantic salmon are distributed across a large latitudinal range and display a wide variety of life history characteristics (Klemetsen et al., 2003), balancing selection on co-adapted linked loci may be important for maintaining adaptive diversity within and among locally adapted populations (Lehnert et al., 2019; Wellband et al., 2019). Indeed, several recent examples have been found within the species, ranging from chromosomal inversions, translocations, fusions, and large haplotype blocks under selection (Lehnert et al., 2019; Wellband et al., 2019; Bertolotti et al., 2020; Stenløkk et al., 2022; Watson et al., 2022; Miettinen et al., 2023).

To date, no studies have identified linked genomic architectures with parallel signal of adaptation across the Atlantic Ocean within the species (but see studies in other species: Bradbury et al., 2010; Nicolas et al., 2025). This may partly be due to large genomic and chromosomal differences between European and North American lineages of Atlantic salmon that have evolved since their divergence over 600,000 years ago (Brenna-Hansen et al., 2012; Lehnert et al., 2020). Nonetheless, despite this divergence, historical trans-Atlantic secondary contact between continents has been documented in certain areas of the range (King et al., 2007), and recent studies have found evidence of introgression of a chromosomal rearrangement in some of these geographic regions (Lehnert et al., 2019; Watson et al., 2022). Therefore, Atlantic salmon is an excellent species to investigate linked genomic architecture, given the richness of structural changes in its genome (Lien et al., 2016; Leitwein et al., 2017; Bertolotti et al., 2020), parallel clines in life history and environment (Metcalfe & Thorpe, 1990; Jeffery et al., 2017), as well as opportunities for trans-oceanic genetic exchange within the species (Lehnert et al., 2019).

In this study, we perform a genome scan to identify large genomic regions that differ significantly from genome-wide population structure for Atlantic salmon. This approach has been previously used to identify large structural variants and haplotype blocks across diverse taxa (Huang et al., 2020; Harringmeyer & Hoekstra, 2022; Dallaire et al., 2024). Using our genomic dataset of over 4,000 individuals from 134 locations spanning both sides of the North Atlantic Ocean (see Figure 1), we identify and focus on a large haplotype block (∼3-Mbp) that is polymorphic across the species range and exhibits clinal patterns in each continent, along with evidence of independent evolutionary origins and introgression. We explore variation this haplotype block, aiming to characterize its structure and role in adaptation. Our work contributes to the growing understanding of the adaptive genomic landscape in Atlantic salmon and provides insight into the role of large-scale linked genomic architectures in salmonid evolution and diversification (Lien et al., 2016), as well as eco-evolutionary processes more broadly.

**Fig. 1.**
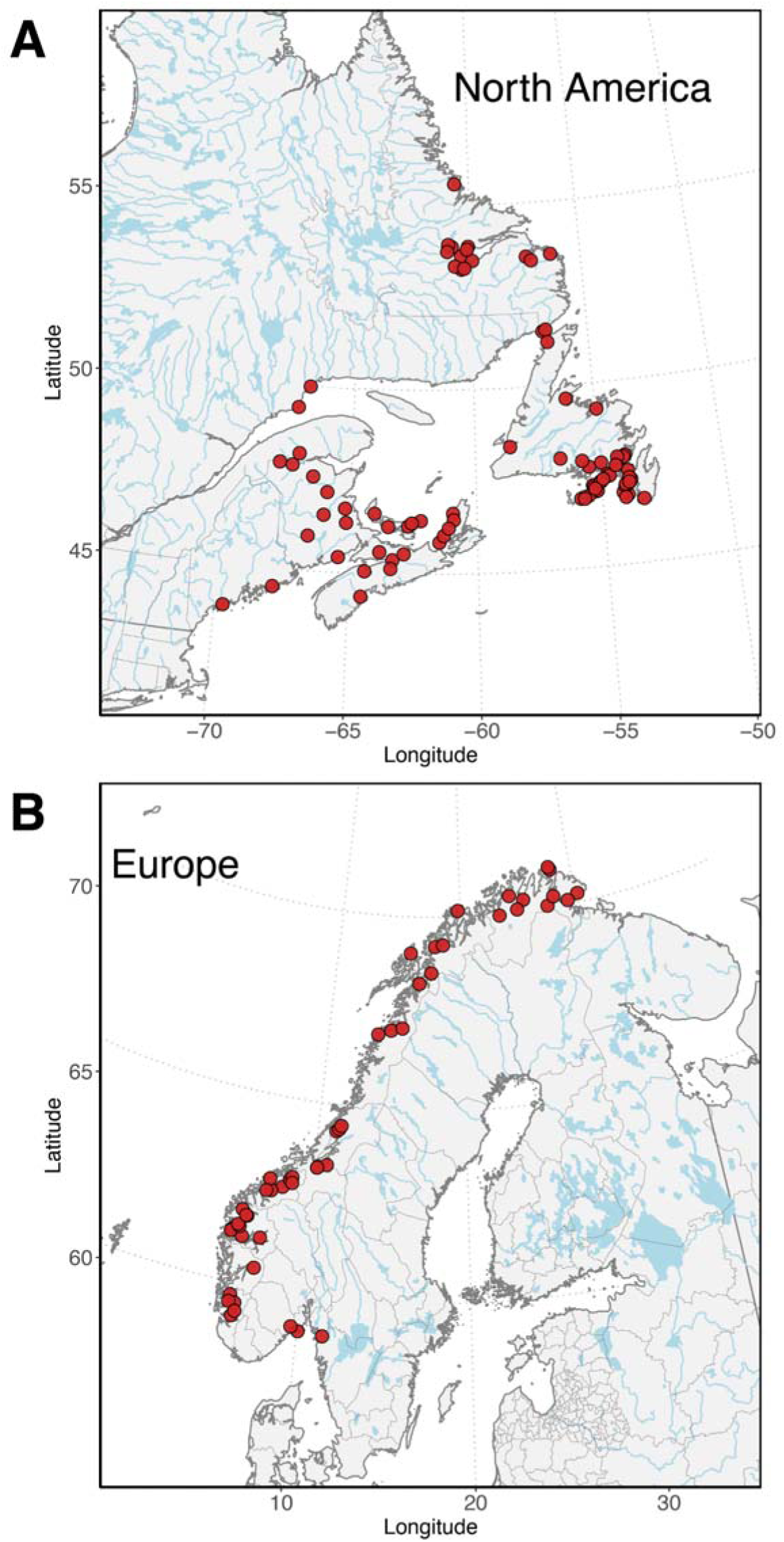
Maps of all Atlantic salmon (*Salmo salar*) sampling sites in (A) North America and (B) Europe that were genotyped with the 220,000 SNP array. Features in maps were generated using ta from https://www.naturalearthdata.com/downloads/ and plotted using the R package *ggplot*.

## Methods

### SNP genotyping

Individuals were genotyped using a targeted bi-allelic SNP Affymetrix Axiom array developed for Atlantic salmon by the Centre for Integrative Genetics (CIGENE, Ås, Norway). The array is comprised of 220,000 SNPs, which are a subset of the 930K XHD *Ssa*l array (dbSNP accession numbers ss1867919552–ss1868858426) (see Barson et al., 2015). Genotype data for individuals from 50 sites in Norway (n=806 individuals) and 84 sites in North America (n= 3456 individuals) were obtained from previous studies (Barson et al., 2015; Sylvester et al., 2018a; Sylvester et al., 2018b; Lehnert et al., 2019; Watson et al., 2022; Nugent et al., 2024) (Table S1). Loci were removed that were deemed low quality for each genotyping run. In total, our full dataset includes 134 Atlantic salmon locations (n = 4262 individuals) spanning from ∼44-56° latitude in North America and ∼56-71° latitude in Europe (Fig. 1). SNPs were filtered for minor allele frequency (MAF) of 0.05 across the range, resulting in 164,579 polymorphic loci with a genotyping rate of 0.997. SNPs are mapped to the newest assembly for Atlantic salmon (Ssal_v3.1) based on the European lineage.

### Local PCAs

To investigate potential linked genomic architecture in the genome, we used local principal component analysis (PCA) using the R package *lostruct* (Li & Ralph, 2019), similar to other recent studies (Huang et al., 2020; Harringmeyer & Hoekstra, 2022). In *lostruct*, we used a window size of 50 SNPs (non-overlapping) and K=2. Each chromosome was analyzed separately, and then multidimensional scaling (MDS) axis 1 values across the genome were compared to identify outlier windows. Outlier windows were those with MDS1 values that were ±3 standard deviations (SD) from the mean, thus corresponding to two-tailed *p*-value <0.0027. Outliers correspond to genomic regions that deviate from the average genome-wide pattern of population structure and thus may represent regions of atypical population structure that may be indicative of divergent haplotype blocks and/or structural variants. MDS axis 2 values were also compared and analyzed; however, we focus on MDS axis 1 for genomic regions that drive major differences relative to backgrounds structure (see Results).

### Linkage disequilibrium and genetic clustering

Using the outlier windows identified by *lostruct*, we then examined linkage disequilibrium (LD). For Europe and North America separately, pairwise LD between all SNPs was calculated using PLINK v1.9 (Purcell et al., 2007) and visualized using *gplots* (Warnes et al., 2016). We next used PCA using the R package *pcadap*t (Luu et al., 2017) to investigate clustering patterns in the outlier regions with K=2 and MAF>0.05. We examined PCAs for clustering patterns that were consistent with large haplotypes and/or structural variants in the genome, where distinct clusters were present that deviated from expected population structure. To assign individuals to clusters in the PCA, we used kmeans function in base R.

### Evidence of trans-oceanic introgression of Ssa18 haplotypes

Our initial genome scan for structural variants identified one novel genomic region of interest on Ssa18 with high LD and evidence of six genotype clusters in PCA (Results). This clustering pattern has been documented in other systems with three haplotypes in regions of reduced recombination, including complex chromosomal rearrangements (González et al., 2014; Knief et al., 2016; Ruiz-Arenas et al., 2019; Ishigohoka et al., 2024), and these haplotypes are herein referred to as A, B, and C (Results).

Haplotypes A and B were common in Europe, whereas A and C were common across the entire range in North America, suggesting a shared ‘ancestral’ A haplotype, with continent-specific alternative haplotypes (B in Europe and C in North America). Yet, based on PCA clustering, the B haplotype (European) was also present in North America but localized to specific geographic regions, which corresponded to known locations of trans-Atlantic secondary contact and introgression in the species (King et al., 2007; Bradbury et al., 2015; Lehnert et al., 2019; Nugent et al., 2024).

Based on PCA clustering patterns, we calculated the frequency of continent-specific alternative haplotypes within each continent. Here, we calculated the frequencies of putative ‘European B allele’ in North American populations and the putative ‘North American C allele’ in European populations and considered this as a measure of potential introgression within populations, allowing us to examine the spatial distribution of Ssa18 haplotype introgression.

### Differentiation among haplotype clusters and continents

We next examined divergence among haplotype clusters (three homozygous genotypes) and continents. Locus-specific *F*_ST_ across Ssa18 was calculated using PLINK v1.9 (Purcell et al., 2007) for various comparisons between continents and between haplotype groups. We focused our comparisons on continental differences within the ‘ancestral’ haplotype groups (AA European individuals vs. AA North American individuals), as well as differences between the ‘ancestral’ AA individuals and individuals of each ‘alternative’ genotype group (BB in Europe and CC in North America) within each continent. Differences in *F*_ST_ within and outside the haplotype block were calculated and statistically compared using a Mann-Whitney U test.

### Evidence of selection and adaptation associated with haplotypes

The results from our above analyses guided the remaining analyses to examine evidence for an adaptive role of the haplotype block, which included testing for evidence of selection on haplotypes, as well as examining associations with geographic, environmental, and life history variation.

First, to investigate evidence of positive selection, we used the Sweepfinder composite likelihood ratio (CLR) method in *SweeD 4.0* (Nielsen et al., 2005; Pavlidis et al., 2013). This method is robust to variation in recombination, ascertainment bias, and demography. For each continent, we separated individuals based on ancestral and continent-specific genotype cluster (AA and BB for European samples; AA and CC for North American samples). CLR values were calculated for Ssa18 using a grid size of 200 windows across the chromosome. We categorized regions with CLR values greater than the 95% quantile of chromosome-wide values as under selection in all comparisons.

Next, for all populations within each continent, we quantified the frequency of the ‘A’ allele (or haplotype) in the population and examined the relationship with latitude. These relationships were evaluated using separate generalized linear models (GLMs), where R^2^ for each model was calculated as the residual deviance divided by the null deviance and subtracted from 1. Evidence of latitudinal associations were apparent (Results), and thus we examined the association between the A allele with climate and life history, as both variables are linked to latitude. For climate data, 19 bioclimatic variables (Fick & Hijmans, 2017) (Table S2) were accessed using the R package *rbioclim* (Exposito-Alonso, 2017) and extracted for each population using latitude and longitude coordinates. For life history, we examined the relationship with smolt age, which is the age that salmon migrate from freshwater to the marine environment. Smolt age is an important life history trait that is heritable (Páez et al., 2011), has an unknown genomic basis, and varies with latitude and environmental condition (Thorpe, 1977; Power, 1981; Metcalfe & Thorpe, 1990). We used the same approach as above (GLM) to quantify the relationship between the frequency of the ‘A’ allele and both bioclimatic variables and smolt age, where we analyzed a reduced dataset for only populations with smolt age information (see Supplement for more details and Table S3).

With a focus on the adaptive role of the haplotype block, we also examined gene annotation within the region. Genes within the Ssa18 haplotype block were accessed from NCBI (RefSeq GCF_905237065.1) based on the newest assembly for Atlantic salmon (Ssal_v3.1).

## Results

### Local PCAs

To investigate large-scale haplotype blocks, we used local PCAs and found seven outlier windows (*p*-value ≤ 0.0027) based on MDS1 values across the genome (Figure 2A). Outlier windows corresponded to genomic regions on four chromosomes, including nearby regions on Ssa08 (spanning 1.32-5.04-Mbp), Ssa23 (spanning 0.59-5.47-Mbp), Ssa18 (44.88-52.52-Mbp), and Ssa12 (11.44-13.65-Mbp). The region on Ssa12 did not show patterns consistent with multiple haplotypes. The regions on Ssa08 and Ssa23 have previously been associated with structural variants (chromosomal rearrangement) in Atlantic salmon (Lehnert et al., 2019; Wellband et al., 2019; Watson et al., 2022), highlighting the strength of this method to identify large-scale linked genomic architectures. However, to date, the region identified on Ssa18 has not been previously known to harbour any structural variants or major haplotype blocks; nevertheless, it emerged as the most significant outlier region in our analysis. We thus focused our analysis on exploring this previously unknown genomic feature. However, we also note that additional outlier regions were identified based on the second MDS axis (MDS2), including the same region on Ssa18, along with 30 other outlier regions (Table S4; Fig. S1). Most outlier regions on MDS2 did not show clear clustering patterns (see Fig. S1) like Ssa18 (see below) and may represent regions of the genome where divergence is reduced between Europe and North American salmon (i.e., deviating from average population structure genome-wide). However, we note that some regions do show clustering patterns that may be consistent with divergent haplotypes, such as inversions or other structural variants that may warrant further investigation (e.g., Ssa26; see Fig. S1).

**Fig. 2.**
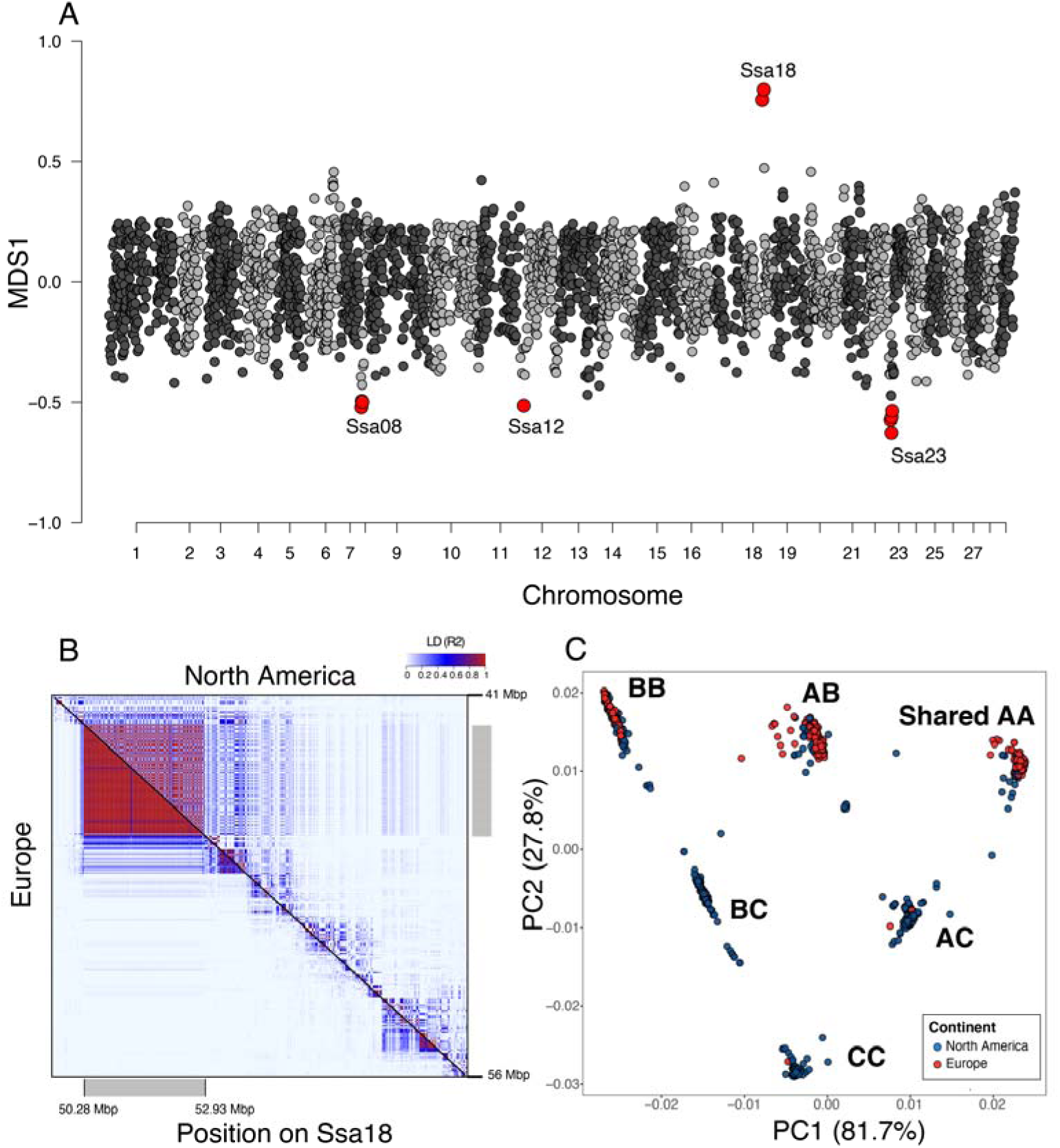
**(A)** Manhattan plot of multidimensional scaling axis 1 (MSD1) values from local principal component analysis (PCA). Red points indicate MDS1 outlier windows (±3 standard deviations from the mean; *p*-value ≤ 0.0027) which may be indicative of structural variants. For the region identified on Ssa18, panel **(B)** shows pairwise linkage disequilibrium (LD; r^2^) between SNPs in the region with European values below diagonal and North American above diagonal. High LD is indicated by red and any fixed loci are indicated by white (LD not calculated). **(C)** Within the region of high LD (gray bar in panel B), genotypes of individuals clustered into six groups using principal component analysis (PCA). Clusters corresponded to genotypes indicated by labels. All analyses were based on 220K SNP data from 134 populations (n=4262 individuals).

### Linkage disequilibrium and genetic clustering

In the region identified on Ssa18, we investigated linkage disequilibrium (LD) and clustering patterns. A 2.65-Mb region of high LD (approximately 50.28-52.93-Mbp), consistent with a large haplotype block, was identified (Fig. 2B). LD was higher in Europe compared to North America. LD was also examined beyond the region identified (30-60-Mbp; see Fig S2), revealing some smaller LD blocks outside of this region. However, we note the absence of SNPs downstream of 44.9-49.6-Mbp in this region. Clustering patterns based on principal component analysis (PCA) revealed six genotype clusters within the region, a pattern previously observed for complex rearrangements with three haplotypes in other species including humans and birds (González et al., 2014; Knief et al., 2016; Ruiz-Arenas et al., 2019). As outlined briefly in the Methods, we identified three clusters that were common in Europe (Fig. S3) consistent with two haplotypes (A and B) and thus three genotypes herein called AA, AB, and BB (Fig. 2C). In North America, six clusters were observed representing three haplotypes (herein A, B, and C) and their possible genotype combinations (Fig. 2C and Fig. S3). Notably, within North America, it appears that the A and C haplotypes are common and present across the entire range, whereas the B haplotype is limited to specific geographic regions (see more below). This suggests that the B haplotype may have been introduced from Europe through secondary historical contact, as indicated by its geographic distribution (King et al., 2007; Lehnert et al., 2019). This is consistent with evidence of introgression of other genomic regions in the Atlantic salmon genome, including structural variants (Lehnert et al., 2019; Watson et al., 2022; Nugent et al., 2024). Therefore, we hypothesize that the A haplotype is the ancestral shared haplotype, with continent-specific alternative haplotypes (B in Europe and C in North America) arising independently in the same genomic region in each lineage following their split over 600,000 years ago (King et al., 2007; Rougemont & Bernatchez, 2018).

### Evidence of trans-oceanic introgression of Ssa18 haplotypes

As indicated, there was potential evidence of trans-oceanic introgression of haplotypes. We evaluated the distribution of continent-specific alleles as a potential signal of introgression within each continent (e.g., ‘European B allele’ in North America and ‘North American C allele’ in Europe). Allele frequencies for each continent are shown in Figure 3A and 3B. Based on the occurrence of these continent specific alleles, we estimated that introgression was present in four populations in northern Norway (1.2% of European individuals; n=10) and 76% of sites in North America (41% of North American individuals, n=1404; see Fig. S4). Notably, the higher number of introgressed populations and individuals in North America is partly due to high sampling coverage in geographic regions of known secondary contact (e.g., southern Newfoundland) (Bradbury et al., 2015; Lehnert et al., 2019; Watson et al., 2022; Nugent et al., 2024).

**Fig. 3.**
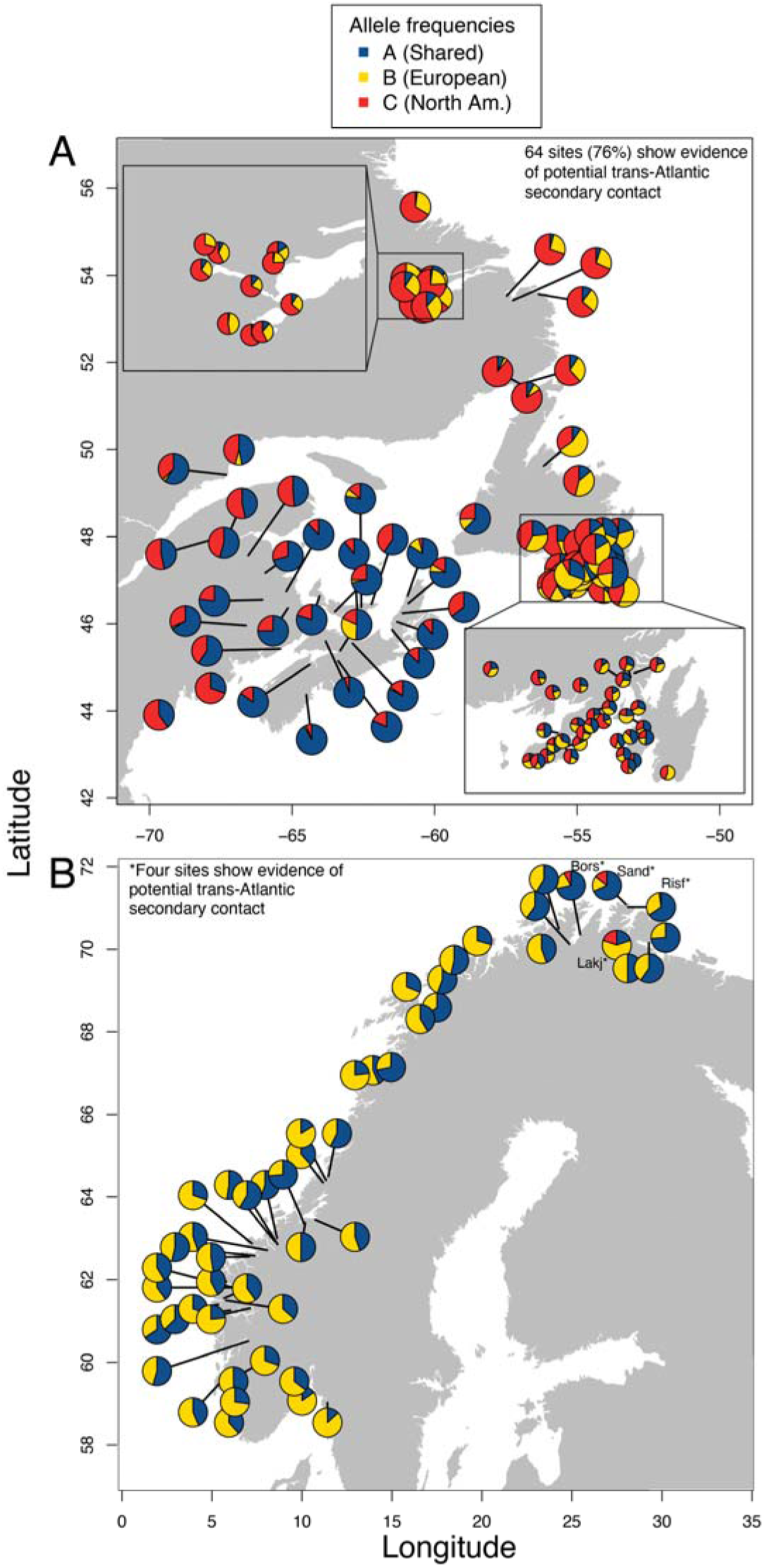
Maps of allele frequencies in **(A)** North America and **(B)** Europe, respectively. Asterisks in B represent populations with evidence of secondary contact; see supplement for populations with contact in panel A.

### Differentiation among haplotype clusters and continents

We measured genetic differentiation (locus-specific *F*_ST_) across Ssa18 for various comparisons (Fig. 4). Between Europe and North America for the shared ancestral genotype group (AA), we found a significant difference in *F*_ST_ values within and outside the haplotype block (Mann-Whitney U test: W =12439, p-value < 2.2e-16). Mean *F*_ST_ was 2.5X greater outside the haplotype block (*F*_ST_ =0.360) compared to within (*F*_ST_ =0.023) (Fig. 4A,D). Notably, approximately 60% of SNPs (n=64 out of 106 SNPs in total) within the haplotype block were fixed for the same allele in both continents; whereas, outside of this region, no loci were fixed for the same allele in both continents (Figure 4A and 4D). Here, we note that loci fixed for the same allele are not included in our estimates of mean *F*_ST_, as *F*_ST_ cannot be calculated for these loci. Differences were also found in the level of divergence between the ancestral shared genotype (AA) and the two continent-specific alternative genotypes (Fig. 4). For instance, for each comparison, *F*_ST_ values were significantly higher within the haplotype block compared to outside of the region. Within the European comparison (AA vs. BB), *F*_ST_ was >100X greater within the haplotype block (*F*_ST_ = 0.989) compared to outside (*F*_ST_ = 0.008) (Mann-Whitney U test: W = 542275, p-value < 2.2e-16). This differed from the North American comparison (AA vs. CC), where *F*_ST_ was approximately 46X greater within the SV (*F*_ST_ =0.814) compared to outside the region (*F*_ST_=0.0175) (Mann-Whitney U test: W =408178, p-value < 2.2e-16). Notably, 72% of SNPs within the region were fixed for different alleles (*F*_ST_ =1.00) in the European genotype comparison (AA vs. BB), whereas in the North American comparison (AA vs. CC), only 25% of loci were fixed for different alleles (Fig. 4), suggesting reduced differentiation between genotype clusters in North America. Similarly, within the North American comparison (AA vs. CC), 36 loci were fixed for the same allele within the haplotype block, whereas no loci were fixed for the same allele in the European comparison (AA vs. BB) (Fig. 4), further highlighting the lower differentiation in North American haplotypes.

**Fig. 4.**
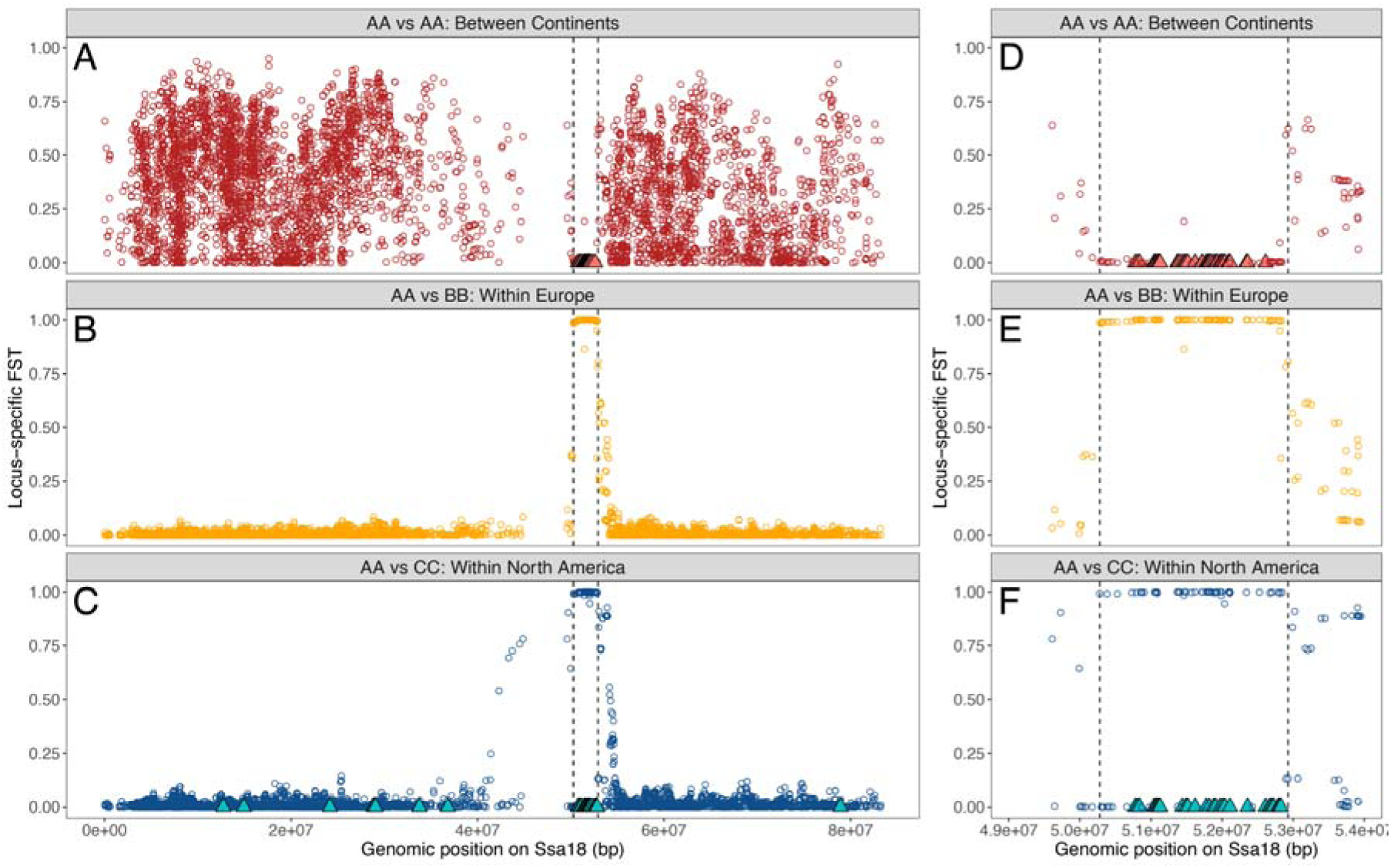
**(A)** Locus-specific *F*_ST_ on Ssa18 between continents for a shared ancestral genotype (AA). Locus-specific *F*_ST_ on Ssa18 *F*_ST_ between the AA genotype and continent-specific alternative genotypes within **(B)** Europe (BB genotype) and within **(C)** North America (CC genotype). Genomic region of interest on Ssa18 is highlighted by dashed lines. Adjacent panels **(D,E,F)** show same comparisons zoomed in on the specific region of interest on Ssa18. Note that *F*_ST_ cannot be calculated for loci that were fixed for the same allele between groups and therefore *F*_ST_=0 are shown as triangles for these loci.

### Evidence of selection and adaptation associated with haplotypes

Evidence of selection on the haplotypes was apparent in both continents when examining signatures of selection on Ssa18 within genotype groups. We used *SweeD 4.0* to calculate composite likelihood ratio (CLR) across chromosome Ssa18 and found significant evidence (>95% quantile) of positive selective sweeps on both genotypes (AA and continent-specific alternative genotypes; BB in Europe and CC in North America) within each continent within the same genomic region of interest (Fig. 5A,B).

**Fig. 5.**
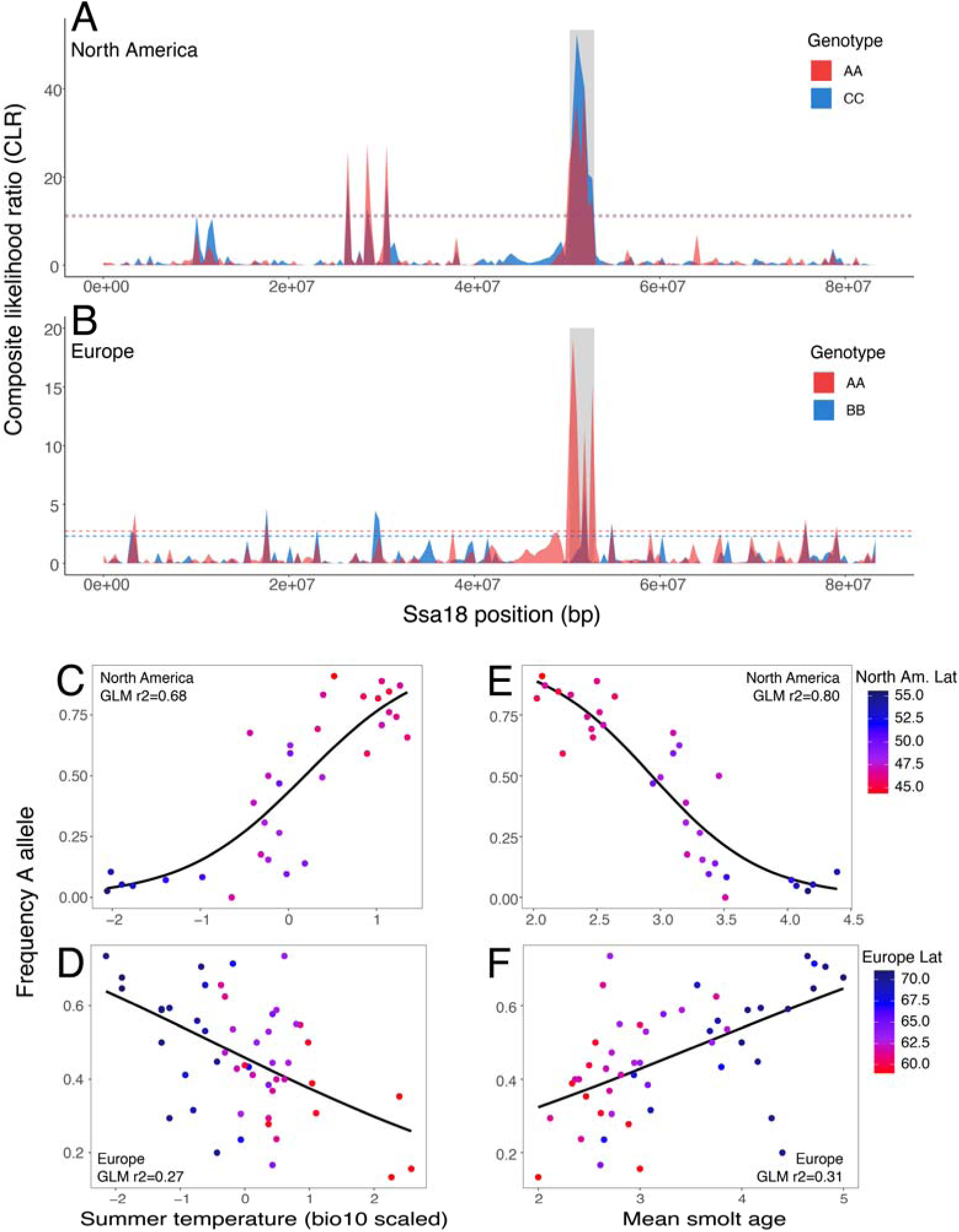
Continent-specific differences associated with smolt age and the haplotype groups in Atlantic salmon (*Salmo salar*). Evidence of selection in **(C)** North America and **(D)** Europe for each genotype (AA in red; continent-specific alternative genotype in blue) are quantified based on composite likelihood ratio (CLR). For the alternative genotype, analyses include only individuals that were North American origin (CC) or European origin (BB) for each continent separately. The dotted lines represent the top 95% quantile of values for each genotype. Relationship for generalized linear model (GLM) between summer temperature (BIO10) and frequency of the A allele in **(C)** North America and **(D)** Europe with the r^2^ for each model provided in the plot. Points for populations are coloured by latitude.

Next, we examined evidence of potential adaptation associated with the putative haplotypes, including by evaluating the relationship between allele frequency and latitude, climate, and life history in both Europe and North America. Latitudinal clinal variation in the frequency of the ancestral A allele was found in both continents, although with differing magnitude and direction of relationships: North America (GLM estimate=-0.069, SE=0.0078, *p*=1.28e-13, r^2^=0.49); Europe (GLM estimate=0.0149, SE=0.0052, *p*=0.006, r^2^=0.15) (Fig. 5C-F; Fig. S5). In both continents, frequencies of the A allele were most strongly associated with summer temperatures (BIO10 mean temperature in the warmest quarter; Fig. 5 C,D) with highly significant but again opposing directions for the relationships (North America: estimate=0.260, SE=0.0319; *t*=8.128, *p*<3.52e-09; r^2^=0.68; Europe: estimate=-0.082, SE=0.0.0197; *t*=-4.183, *p*=0.0001; r^2^=0.27). Relationships between allele frequencies and other environmental variables are provided in the Supplement (Fig. S6-7, and Table S2).

Similarly, mean smolt ages were highly correlated with the frequencies of the A allele on both continents; however, the relationship was in opposing directions (Fig. 5E and 5F and Fig. S5). The strength of the relationship between smolt ages and allele frequencies was two-fold higher in North America (GLM estimate=-0.419, SE=0.0378, *p*=2.568e-12, r^2^=0.80) relative to Europe (GLM estimate=0.110, SE=0.0238, *p*=2.99e-5, r^2^=0.31). Therefore, despite the A allele being hypothesized as ancestral and shared between continents, our results suggest the A allele is associated with younger smolts in North America but older smolts in Europe (see Fig. 5E and 5F). Given these associations, we also examined gene annotation within the Ssa18 regions. A list of genes within the genomic region are provided in Table S5.

### Characterizing the genome structure of haplotype block using long-read data

Based on the results reported here, we characterized the genomic structure of the haplotype block. Long-read sequencing data were evaluated from individuals from both Newfoundland and Norway. This included two individuals from Garnish River, Newfoundland (GCA_931345325.1, GCA_923944775.2), which were characterized as having CC and BC genotypes based on SNP array data. In addition, salmon from geographic regions in Norway (GCA_931345835.1, GCA_931347365.1, GCA_931345645.1, GCA_931345955.1, GCA_931345925.1, GCA_931346935.2 and GCA_905237065.2) that would be expected to have common European genotypes (AA, AB, BB) were also included. Individuals were sequenced using the PromethION instrument from Oxford Nanopore Technologies, and assemblies were polished using Illumina reads (described in Stenløkk et al., 2022). Despite the expected differences in haplotypes, dot plot visualizations of genome-genome alignments did not exhibit any discontinuity patterns in sequence order characteristic of inversions or other large structural variants.

## Discussion

Advances in sequencing technology are enabling the discovery of linked genomic architectures, such as large haplotype blocks and structural variants, that can contribute to adaptation (Huang et al., 2020; Mérot et al., 2020; Todesco et al., 2020; Harringmeyer & Hoekstra, 2022; Rubin et al., 2022; Battlay et al., 2023). Here, we identify a novel and complex haplotype block on chromosome Ssa18, that is putatively under selection and strongly associated with latitude, climate, and life history (smolt age) in divergent lineages of Atlantic salmon across the Atlantic Ocean. Clustering patterns were consistent with three haplotypes in a species-wide region of low recombination (Ishigohoka et al., 2024), with patterns consistent with those detected for complex chromosomal rearrangement in other species (González et al., 2014; Knief et al., 2016; Ruiz-Arenas et al., 2019). Data support the inference that haplotype ‘A’ is ancestral and shared between allopatric lineages of Atlantic salmon. Haplotypes divergent from A likely evolved separately within each lineage after their divergence, yet there was evidence of introgression of these divergent haplotypes within each lineage, consistent with historical trans-oceanic secondary contact (King et al., 2007; Bradbury et al., 2015; Rougemont & Bernatchez, 2018; Lehnert et al., 2019; Nugent et al., 2024).

Our study provides the first evidence of large-scale haplotype block that is putatively adaptive and polymorphic across divergent lineages of Atlantic salmon, offering novel insight to the species’ evolutionary history. In European populations, we primarily detected three genotype clusters (AA, AB, BB; see Fig. S3), consistent with clustering patterns often observed for inversions (Ruiz-Arenas et al., 2019; Harringmeyer & Hoekstra, 2022). In North American populations, three genotype clusters were also common across the range (AA, AC, CC), with one of these (AA) shared with Europe. These results support a common ancestral haplotype (‘A’), and the independent origins of divergent haplotypes (‘B’ and ‘C’) within the same Ssa18 region in each continent. Notably, comparisons of differentiation among haplotype clusters within continents supported lower differentiation within the North American comparison (AA vs. CC) relative to the European comparison (AA vs. BB). Reduced differentiation within the North American comparison may be attributed to generally lower diversity and reduced population structure in this lineage, likely resulting from different postglacial colonization histories (King et al., 2001; Bourret et al., 2013). This further supports the hypothesis that the haplotypes arose independently in both lineages after their divergence. Alternatively, though less likely, differences in diversity could be attributed to ascertainment bias, as the SNP array was originally developed using Atlantic salmon of European origin (Barson et al., 2015).

While three genotypes were common across the sampling range within each continent, in North America, three additional clusters were abundant and closely related to European clusters (Fig. 2C), suggesting potential for introgression of the European haplotype. PCA clustering patterns were also consistent with introgression in geographic regions in North America, including Newfoundland and Labrador, where trans-oceanic secondary contact has been documented by previous studies from mitochondrial and genomic datasets (King et al., 2007; Bradbury et al., 2015; Lehnert et al., 2019; Watson et al., 2022). This includes recent work that found introgression of a European chromosomal rearrangement between Ssa01 and Ssa23 within North American populations (Lehnert et al., 2019; Watson et al., 2022), further supporting a role for adaptive introgression of linked genomic architecture. In addition, clustering patterns supported evidence of trans-oceanic introgression in four populations in northern Europe, consistent with studies using a small number of nuclear and mitochondrial markers in northern populations from the Barents Sea (Asplund et al., 2004; Ryynänen & Primmer, 2004; Makhrov et al., 2005).

The maintenance of divergent haplotypes, along with evidence of introgression, supports the hypothesis of selection acting on the Ssa18 haplotypes. Our analyses also revealed signals of positive selection acting on haplotype groups within each continent. Moreover, we found a strong relationship between Ssa18 genotype and life history (smolt age) at the population level, a trait for which a genomic basis has not yet been identified. In Atlantic salmon, the age of smoltification can vary from one year to eight years (Klemetsen et al., 2003). It generally increases with latitude and can be dependent on environmental factors that influence growth potential (Thorpe, 1977; Metcalfe & Thorpe, 1990). Thus, smolt age is expected to be linked to freshwater growth opportunity, which is maximized in summer months (Power, 1981; Thorpe et al., 1989; Metcalfe & Thorpe, 1990). We also found a strong association between Ssa18 genotype and summer temperature. While environment clearly plays a role in smolt age, studies suggest that the heritability of smolt age is approximately three times higher in North America relative to Europe (Refstie et al., 1977; Páez et al., 2011). This is consistent with a stronger association between A haplotype frequency and phenotype in North American (R^2^=0.80) than in European (R^2^=0.31) populations in our study. Further, several genes within the Ssa18 region (see Table S5) may play a role in the smoltification process, further highlighting a role in adaptation. For example, the region harbours genes that have been linked to migration (*protocadherin-7-like*) (Lemopoulos et al., 2018), metamorphosis (*inhibitor of growth protein 2-like*; *ING2-l*) (Wagner et al., 2001), and photoreception, olfaction, or skin pigmentation (*Sodium/potassium/calcium exchanger 2-like*) (Schnetkamp, 2013).

Admittedly, while the above evidence linking Ssa18 genotype and smolt age is compelling, population-level associations showed opposing relationships in each continent with different genetic effects. While perplexing, the opposing relationships between continents may reflect the large differences between the alternative haplotypes (‘B’ and ‘C’) which may have differing effects on phenotype. While epistasis or environment interactions may also explain these results, it is also possible that this genomic region does not directly influence life history but may instead be physically linked to loci that do. This may explain differing parallel but inverse relationships on each side of the Atlantic, particularly if this region is more susceptible to the effects of genetic hitchhiking due to an expected lack of recombination in the region (Barton, 2000). Therefore, given that salmon populations recolonized rivers from the south to the north following the last glaciation (King et al., 2001), the association of genotype with latitude and population structure, as well as other factors related to these (environment and life history), may be an artefact of colonization history. Regardless of the mechanisms, this genomic region provides a means to predict population level characteristics across a large geographic range (Lotterhos, 2023), and may be linked to other important traits, and common garden experiments would be beneficial to fully understand the role of the Ssa18 region in driving adaptation (Mérot et al., 2020).

Similarly, the potential association between Ssa18 genotype and other life history traits, particularly those with latitudinal clines (e.g., timing of migration, precocial maturation) (Valiente et al., 2005; Dempson et al., 2017; Vollset et al., 2021), warrant further investigation. For example, recent findings by Beck et al. (in prep.) provide further evidence supporting the potential (mal)adaptive role of Ssa18 in Atlantic salmon life-history traits. Genetic variation in this region was strongly correlated with late migration timing at a population level in North American Atlantic salmon, with a higher prevalence of one genotype occurring in populations from the Maritimes – which have a later migration timing – compared to those in Newfoundland and Labrador.

Many studies have identified similarly large haplotype blocks in salmonids and other species (Stenløkk et al., 2022), however, the majority of those studies often confirm or infer the presence of structural variants, like chromosomal rearrangements, as a mechanism maintaining haplotypes. Despite the large size of the Ssa18 haplotype block (∼3-Mbp), long read sequencing data did not confirm the presence of a structural variant, such as an inversion, suggesting the evolution of co-adapted alleles without structural change. While we do not have data to support a hypothesis for the formation of these haplotypes at this time, the haplotype block is located within the fused region of Ssa18 (Lien et al., 2016), and may retain signals of an ancestral fusion polymorphism that has since become fixed. Haplotypes may be maintained through reduced recombination in this region due to high similarity with homoeologous chromosomes (Lien et al., 2011) and selection. Although we cannot confirm the nature of or mechanism maintaining this large haplotype block, our study highlights the power of the SNP dataset to identify large-scale linked genomic architectures, as our initial genome scan using local PCAs also detected known chromosomal rearrangements on Ssa08 and Ssa23 that have been linked to environmental adaptation (Wellband et al., 2019; Watson et al., 2022). Additional genomic regions identified here using local PCAs (see Fig. S1), as well as in other studies (Bertolotti et al., 2020; Stenløkk et al., 2022), highlight important axes of genomic diversity that remains to be understood.

In conclusion, our study reports the first evidence of a large haplotype block in Atlantic salmon that is polymorphic across the range, as well as strongly linked to climate and life history on both sides of the Atlantic Ocean. Although long-read sequencing data did not reveal a structural mechanism driving haplotype differences, our work provides support that this region is complex, likely not neutral, and may underpin (or be linked) to important adaptation. While more remains to be understood about Ssa18, the genotype-environment associations identified here can still be helpful for predicting population adaptation as well as forecasting future changes (Lotterhos, 2022). Our study contributes to understanding the role that linked architectures play in evolutionary processes, as well as to the ongoing characterization of the adaptive genomic landscape in the Atlantic salmon serving to better understand their role in salmonid evolution and diversification (Lien et al., 2016).

## Acknowledgments

The data used in this study were generated from previous work. Nonetheless, the authors thank the staff of the Newfoundland DFO Salmonids section for juvenile sampling, the Aquatic Biotechnology Laboratory of the Bedford Institute of Oceanography for sample preparation, and CIGENE for SNP genotyping and data processing. We also thank MFFP for collection of samples in the Quebec region. We also thank the Nunatsiavut Government, the Sivunivut Inuit Community Corporation, the Innu Nation, the Labrador Hunting and Fishing Association and local fishers for their support and active participation in sample collection in the Labrador region. This work was supported by Fisheries and Oceans Canada’s (DFO) Programme for Aquaculture Regulatory Research (PARR) and NSERC.

## Funding

This research was supported by NSERC and Fisheries and Oceans Canada.

## Author contributions

Sarah J. Lehnert, Ian R. Bradbury, Sigbjørn Lien, and Tony Kess contributed to the conception and design of the study. Ian R. Bradbury, Sigbjørn Lien, and Nicola Barson provided molecular data and metadata for the study. Sigbjørn Lien and Sarah J. Lehnert performed statistical analyses. Sarah J. Lehnert drafted the manuscript and all authors contributed to the writing and approved the final draft of the manuscript.

## Competing interests

Authors declare no competing interests.

## Data and materials availability

Data for this project were compiled from datasets of previous studies (Barson et al., 2015; Sylvester et al., 2018a; Sylvester et al., 2018b; Lehnert et al., 2019; Watson et al., 2022; Nugent et al., 2024). All scripts used for analyses are available at: www.github.com/SarahLehnert/AtlanticSalmon_Ssa18/.

## Supplementary Materials

Supplemental Methods

Figures S1-7

Tables S1-S5

## Notes

### Competing Interest Statement

The authors have declared no competing interest.

