## Supplemental for "Large haplotypes linked to climate and life history variation in divergent lineages of Atlantic salmon (*Salmo salar*)"

**This PDF file includes:**

Supplemental methods

Tables S1 to S5

Figs. S1 to S7

Supplemental references

Supplemental Methods

Life history: Population level smolt age information

Smolt age information was used from population level data for North America (Table S3), whereas, we used individual-level phenotype data to calculate mean smolt age for each population in Europe (data from Barson et al., 2015). In this case, variation across populations may be non-random in Europe because samples were collected to represent variation in age-at-maturity (Barson et al., 2015). For North America, mean smolt age were compiled (Table S3) from previous work (Hutchings & Jones, 1998; Chaput et al., 2006; Dauphin, 2022; April et al., 2023; Cairns et al., 2023; Douglas et al., 2023; Reader et al., 2024; Kelly et al., in press-a, in press-b) for 33 locations, where mean age data were often available for large and small salmon categories combined. For any rivers with data for large and small salmon separately in (Chaput et al., 2006), averages for both large and small were used to calculate mean smolt age if both were available. For two river systems (Margaree and Restigouche), genetic samples from different tributaries of the rivers were combined (see Table S3), but for Miramichi where data on smolt age were available for both the Northwest and Southwest branches of the river system separately, and thus both sites were included. Additionally, we excluded sites from Maine, USA from the analysis because these populations have been heavily stocked and therefore hatchery practices may result in deviations of smolt age relative to the wild population.

**Table S1.** Site locations for North American and European Atlantic salmon (*Salmo salar*) populations that have been genotyped with the 220,000 SNP array. Genotype data were obtained from previous studies (Barson et al., 2015; Sylvester et al., 2018a; Sylvester et al., 2018b; Lehnert et al., 2019; Watson et al., 2022) as well as additional samples genotyped using the same technology. Locations are organized alphabetically by population code.

| **Site Name** | **Code** | **Lat** | **Long** | **n** | **Continent** |
| --- | --- | --- | --- | --- | --- |
| Riviere Aux Rochers | ARO | 50.00 | -66.86 | 48 | North America |
| Bay de L'Eau River | BDL | 47.51 | -54.73 | 91 | North America |
| Bay du Nord | BDN | 47.73 | -55.44 | 20 | North America |
| Black River | BLA | 47.89 | -54.17 | 82 | North America |
| Branch River | BRA | 46.89 | -53.97 | 68 | North America |
| Big Salmonier Brook | BSA | 47.06 | -55.22 | 81 | North America |
| Big Salmon | BSR | 45.42 | -65.41 | 22 | North America |
| Cape Caribou | CB | 53.62 | -60.42 | 21 | North America |
| Come By Chance River | CBC | 47.97 | -53.96 | 61 | North America |
| Cheticamp River | CHT | 46.64 | -60.95 | 12 | North America |
| Caroline River | CL | 53.25 | -60.42 | 18 | North America |
| Campbellton | CMP | 49.28 | -54.93 | 25 | North America |
| Conne | CNR | 47.91 | -55.70 | 209 | North America |
| Crooked River | CR | 53.87 | -60.83 | 21 | North America |
| Cape Roger Brook | CRB | 47.44 | -54.69 | 56 | North America |
| Cuslett Brook | CUS | 46.96 | -54.16 | 87 | North America |
| Dollards Brook | DLR | 48.02 | -56.57 | 26 | North America |
| Eagle River | EA | 53.53 | -57.47 | 21 | North America |
| Fair Haven Brook | FHB | 47.54 | -53.89 | 103 | North America |
| Flat Bay Brook | FLB | 48.41 | -58.58 | 24 | North America |
| Forteau River | FO | 51.48 | -56.94 | 21 | North America |
| Gaspereau River | GAK | 45.06 | -64.38 | 26 | North America |
| Garnish | GAR | 47.23 | -55.35 | 22 | North America |
| Great Barasway Brook | GBW | 47.12 | -54.06 | 89 | North America |
| Great Rattling Brook - Exploits | GRB | 49.62 | -56.17 | 26 | North America |
| Hunt River | HU | 55.57 | -60.67 | 19 | North America |
| Graham River | JGC | 45.86 | -61.49 | 11 | North America |
| Kenamu River | KE | 53.48 | -59.91 | 22 | North America |
| Kedgwick | KED | 47.91 | -67.91 | 15 | North America |
| Kouchibouguac | KOU | 46.74 | -65.20 | 31 | North America |
| LaHave | LAH | 44.37 | -64.50 | 22 | North America |
| Lance River | LAN | 46.82 | -54.07 | 9 | North America |
| Little Barasway Brook | LBB | 47.18 | -54.03 | 15 | North America |
| Long Harbour | LHR | 47.82 | -54.94 | 18 | North America |
| L'anse au Loup River | LL | 51.53 | -56.82 | 22 | North America |
| Little Salmonier | LSR | 47.07 | -55.18 | 17 | North America |
| Lawn River | LWN | 46.95 | -55.54 | 80 | North America |
| Mabou River | MAB | 46.04 | -61.31 | 27 | North America |
| Matapedia | MAT | 48.18 | -67.14 | 15 | North America |
| Northeast Margaree | MNE | 46.47 | -60.92 | 12 | North America |
| Morells | MOR | 46.30 | -62.71 | 18 | North America |
| Southwest Margaree | MRS | 46.24 | -61.12 | 14 | North America |
| Miramichi-Upper Southwest | MSW | 46.55 | -66.04 | 23 | North America |
| Mulligan River | MU | 53.87 | -60.09 | 18 | North America |
| Miramichi-Upper Northwest | MUN | 47.17 | -65.94 | 24 | North America |
| North Brook Trepassey | NBT | 46.74 | -53.36 | 25 | North America |
| Northeast Complex-1 (PEI) | NEP | 46.45 | -62.21 | 27 | North America |
| Northeast Complex-2 (PEI) | NET | 46.38 | -62.57 | 24 | North America |
| Narraguagus River | NGR | 44.52 | -67.86 | 21 | North America |
| North Harbour River | NHR | 47.92 | -54.03 | 117 | North America |
| Northwest Brook (Mortier Bay) | NMB | 47.17 | -55.32 | 87 | North America |
| Nonsuch River | NON | 47.45 | -54.64 | 91 | North America |
| Northeast Placentia River | NPR | 47.29 | -53.80 | 81 | North America |
| North River NS | NRH | 45.38 | -63.31 | 22 | North America |
| Nashwaak | NSH | 45.96 | -66.62 | 19 | North America |
| Northwest Complex(PEI) | NWP | 46.63 | -64.04 | 17 | North America |
| Paradise River | PA | 53.42 | -57.25 | 19 | North America |
| Patapedia | PAT | 47.86 | -67.39 | 24 | North America |
| Piercey's Brook | PBR | 46.88 | -55.86 | 83 | North America |
| Pipers Hole River | PHR | 47.93 | -54.27 | 87 | North America |
| East River Pictou | PIE | 45.54 | -62.88 | 23 | North America |
| Peters River | PR | 53.34 | -60.71 | 21 | North America |
| Red Harbour River East | RHA | 47.33 | -54.99 | 91 | North America |
| Red Harbour River West | RHW | 47.30 | -55.02 | 75 | North America |
| Richibucto | RIC | 46.36 | -65.15 | 31 | North America |
| River Philip | RPH | 45.59 | -63.82 | 17 | North America |
| Rushoon River | RUS | 47.37 | -54.92 | 84 | North America |
| Red Wine River | RW | 53.93 | -61.00 | 22 | North America |
| South Central PEI | SCP | 46.28 | -63.49 | 14 | North America |
| Sand Hill River | SH | 53.57 | -56.35 | 19 | North America |
| Sandy Harbour River | SHA/SHR | 47.71 | -54.36 | 74 | North America |
| Ship Harbour Brook | SHI | 47.35 | -53.87 | 82 | North America |
| Sheepscot River | SHP | 43.91 | -69.67 | 20 | North America |
| Sebaskachu River | SK | 53.79 | -60.14 | 22 | North America |
| Southeast Placentia River | SPR | 47.23 | -53.88 | 96 | North America |
| Susan River | SR | 53.74 | -61.04 | 22 | North America |
| Stewiacke | STW | 45.14 | -63.38 | 22 | North America |
| Taylor Bay Brook (Burin Penn) | TBR | 46.88 | -55.71 | 80 | North America |
| Tides Brook | TDS | 47.13 | -55.26 | 68 | North America |
| Traverspine River | TR | 53.28 | -60.28 | 22 | North America |
| Riviere de la Trinite | TRI | 49.42 | -67.30 | 49 | North America |
| Upsalquitch | UPS | 47.57 | -66.54 | 28 | North America |
| Western Arm | WAB | 51.19 | -56.76 | 18 | North America |
| Altaelva | Alta | 69.97 | 23.37 | 19 | Europe |
| Årdalselva | Arda | 59.14 | 6.17 | 18 | Europe |
| Årgårdsvassdraget | Arga | 64.31 | 11.22 | 13 | Europe |
| Arøyelva | Aroy | 61.27 | 7.17 | 16 | Europe |
| Aursunda | Aurs | 64.37 | 11.39 | 18 | Europe |
| Beiarvassdraget | Beia | 67.03 | 14.58 | 15 | Europe |
| Børselva in Porsanger | Bors | 70.31 | 25.52 | 17 | Europe |
| Daleelva Høyangervassdraget | Dale | 61.22 | 6.07 | 12 | Europe |
| Dalselva in Dale | Dals | 61.36 | 5.40 | 17 | Europe |
| Eidfjordvassdraget | Eidf | 60.47 | 7.07 | 21 | Europe |
| Eira | Eira | 62.68 | 8.12 | 18 | Europe |
| Elvegårdselva (Bjerkvik) | Elve | 68.55 | 17.56 | 16 | Europe |
| Enningdalselva | Enni | 58.98 | 11.47 | 15 | Europe |
| Etneelva | Etne | 59.67 | 5.93 | 16 | Europe |
| Flekkeelva | Flek | 61.31 | 5.35 | 19 | Europe |
| Forsåvassdraget | Fors | 68.27 | 16.63 | 17 | Europe |
| Gaula in Sør-Trøndelag | GauST | 63.34 | 10.24 | 24 | Europe |
| Gloppenelva | Glop | 61.77 | 6.20 | 25 | Europe |
| Hjalma | Hjal | 61.91 | 5.85 | 7 | Europe |
| Homla | Homl | 63.41 | 10.80 | 18 | Europe |
| Jølstra | Jols | 61.46 | 5.83 | 19 | Europe |
| Komagelva | Koma | 70.24 | 30.25 | 17 | Europe |
| Lakselva in Porsanger | Laks | 70.08 | 24.92 | 16 | Europe |
| Laksjohka | Lakj | 70.06 | 27.55 | 5 | Europe |
| Laukhellevassdraget | Lauk | 69.23 | 17.86 | 17 | Europe |
| Målselvvassdraget | Mals | 69.27 | 18.51 | 16 | Europe |
| Måna | Mana | 62.54 | 7.44 | 14 | Europe |
| Maskejohka | Mask | 70.28 | 28.15 | 1 | Europe |
| Namsen | Nams | 64.46 | 11.52 | 13 | Europe |
| Nausta | Naus | 61.51 | 5.72 | 10 | Europe |
| Numedalslågen | Nume | 59.03 | 10.06 | 16 | Europe |
| Reipåga | Reip | 66.91 | 13.63 | 17 | Europe |
| Repparfjordelva | Repp | 70.45 | 24.32 | 17 | Europe |
| Risfjordvassdraget | Risf | 70.98 | 28.17 | 17 | Europe |
| Roksadalsvassdraget | Roks | 69.05 | 15.87 | 19 | Europe |
| Ryggelva | Rygg | 61.78 | 6.13 | 17 | Europe |
| Saltdalsvassdraget | Salt | 67.10 | 15.42 | 7 | Europe |
| Sandfjordelva in Gamvik | Sand | 71.05 | 28.05 | 17 | Europe |
| Skienselva | Skie | 59.13 | 9.63 | 17 | Europe |
| Skipsfjordvassdraget | Skip | 70.16 | 19.80 | 17 | Europe |
| Søya | Soya | 62.89 | 8.54 | 17 | Europe |
| Suldalslågen | Suld | 59.48 | 6.25 | 16 | Europe |
| Suma | Surn | 62.97 | 8.67 | 20 | Europe |
| Sylteelva in Fraena | Sylt | 62.84 | 7.27 | 18 | Europe |
| Todalselva (Toaa) | Toda | 62.82 | 8.70 | 17 | Europe |
| Tressa | Tres | 62.52 | 7.13 | 18 | Europe |
| Vestre Jakobselve | VeJa | 70.11 | 29.33 | 22 | Europe |
| Vigda | Vigd | 63.31 | 10.18 | 17 | Europe |
| Vikedalselva | Vike | 59.49 | 5.90 | 13 | Europe |
| Vorma | Vorm | 59.27 | 6.33 | 18 | Europe |

Table S2. Bioclimatic variables and strength of association (r^2^ of generalized linear model; GLM) with allele frequency in European and North American Atlantic salmon (*Salmo salar*). For visualization of relationships see Fig. S6 and S7.

|  | **GLM r^2^** | |
| --- | --- | --- |
| **Bioclimatic variables** | **North America** | **Europe** |
| BIO1 = Annual Mean Temperature | 0.411 | 0.187 |
| BIO2 = Mean Diurnal Range (Mean of monthly (max temp - min temp)) | 0.239 | 0.119 |
| BIO3 = Isothermality (BIO2/BIO7) (* 100) | 0.146 | 0.071 |
| BIO4 = Temperature Seasonality (standard deviation *100) | 0.113 | 0.015 |
| BIO5 = Max Temperature of Warmest Month | 0.587 | 0.148 |
| BIO6 = Min Temperature of Coldest Month | 0.023 | 0.105 |
| BIO7 = Temperature Annual Range (BIO5-BIO6) | 0.121 | 0.037 |
| BIO8 = Mean Temperature of Wettest Quarter | 0.100 | 0.008 |
| BIO9 = Mean Temperature of Driest Quarter | 0.098 | 0.056 |
| BIO10 = Mean Temperature of Warmest Quarter | 0.681 | 0.267 |
| BIO11 = Mean Temperature of Coldest Quarter | 0.053 | 0.108 |
| BIO12 = Annual Precipitation | 0.052 | 0.109 |
| BIO13 = Precipitation of Wettest Month | 0.146 | 0.112 |
| BIO14 = Precipitation of Driest Month | 0.082 | 0.122 |
| BIO15 = Precipitation Seasonality (Coefficient of Variation) | 0.146 | 0.019 |
| BIO16 = Precipitation of Wettest Quarter | 0.157 | 0.103 |
| BIO17 = Precipitation of Driest Quarter | 0.032 | 0.118 |
| BIO18 = Precipitation of Warmest Quarter | 0.026 | 0.112 |
| BIO19 = Precipitation of Coldest Quarter | 0.111 | 0.082 |

Table S3. Mean smolt age for North American Atlantic salmon (Salmo salar) for populations that have been genotyped with the 220,000 SNP array. Mean smolt age were accessed from several sources as indicated by references (Hutchings & Jones, 1998; Chaput et al., 2006; Dauphin, 2022; April et al., 2023; Cairns et al., 2023; Douglas et al., 2023; Reader et al., 2024; Kelly et al., in press-a, in press-b). Mean age was determined either from large and/or small salmon in the river. For two river systems (Margaree and Restigouche), samples from different branches of the river were combined (see Codes). Sites from Maine, USA were excluded as these populations have been heavily stocked and the influence of hatchery rearing on smolt age is unknown.

| **River** | **Region** | **Code** | **Mean smolt age** | **Reference** |
| --- | --- | --- | --- | --- |
| Lahave | Maritime | LAH | 2.068 | (Chaput et al., 2006) |
| Gaspereau | Maritime | GAK | 2.196 | (Reader et al., 2024) |
| Kouchibouguac | Gulf | KOU | 2.09 | (Chaput et al., 2006) |
| Cheticamp | Gulf | CHT | 2.294 | (Chaput et al., 2006) |
| Stewiacke | Maritime | STW | 2.027 | (Reader et al., 2024) |
| Richibucto | Gulf | RIC | 2.423 | (Chaput et al., 2006) |
| Margaree | Gulf | MRS & MNE | 2.457 | (Chaput et al., 2006) |
| Morelle | Gulf | MOR | 2.5 | (Cairns et al., 2023) |
| Nashwaak | Maritime | NSH | 2.468 | (Chaput et al., 2006) |
| Big Salmon | Maritime | BSR | 2.23 | (Reader et al., 2024) |
| Miramichi – Southwest | Gulf | MSW | 2.52 | (Douglas et al., 2023) |
| Miramichi- Northwest | Gulf | MUN | 2.55 | (Douglas et al., 2023) |
| East River (Pictou) | Gulf | PIE | 2.64 | (Chaput et al., 2006) |
| Aux Rochers | Quebec | ARO | 2.94 | (April et al., 2023) |
| Restigouche | Gulf | UPS, KED, PAT & MAT | 3.0 | (Dauphin, 2022) |
| Northeast Placentia | Newfoundland | NPR | 3.2 | (Kelly et al., in press-a) |
| De la Trinite | Quebec | TRI | 3.1 | (April et al., 2023) |
| Branch | Newfoundland | BRA | 3.1 | (Hutchings & Jones, 1998) |
| Flat Bay Brook | Newfoundland | FLB | 3.148 | (Chaput et al., 2006) |
| Conne River | Newfoundland | CNR | 3.31 | (Kelly et al., in press-a) |
| Garnish | Newfoundland | GAR | 3.46 | (Kelly et al., in press-a) |
| North Harbour | Newfoundland | NHR | 3.2 | (Hutchings & Jones, 1998) |
| Little Salmonier River | Newfoundland | LSR | 3.21 | (Chaput et al., 2006) |
| Pipers Hole River | Newfoundland | PHR | 3.33 | (Chaput et al., 2006) |
| Exploits River | Newfoundland | GRB | 3.38 | (Kelly et al., in press-a) |
| Campbellton | Newfoundland | CMP | 3.43 | (Kelly et al., in press-a) |
| Northeast Brook Trepassy | Newfoundland | NBT | 3.51 | (Kelly et al., in press-a) |
| Western Arm Brook | Newfoundland | WAB | 3.52 | (Kelly et al., in press-a) |
| Eagle River | Labrador | EA | 4.07 | (Kelly et al., in press-b) |
| Forteau Brook | Labrador | FO | 4.03 | (Chaput et al., 2006) |
| Hunt River | Labrador | HU | 4.16 | (Kelly et al., in press-b) |
| Paradise | Labrador | PA | 4.2 | (Kelly et al., in press-b) |
| Sand Hill River | Labrador | SH | 4.39 | (Kelly et al., in press-b) |

Table S4. Outlier regions (chromosome and genomic position) identified by *lostruct* based on multidimensional scaling (MDS) axis 2 values. Clustering patterns within these regions are shown in Figure S1. Genomic positions are based on the original assembly for Atlantic salmon (Ssal_v3.1). These regions include all outlier windows (50 SNPs) that were directly adjacent to each other.

| **Chr** | **Start (bp)** | **End (bp)** |
| --- | --- | --- |
| 1 | 165148777 | 165447737 |
| 4 | 78366563 | 79058293 |
| 5 | 44018836 | 48386499 |
| 6 | 41647990 | 42077163 |
| 6 | 44357569 | 44573462 |
| 7 | 45773532 | 46039053 |
| 8 | 13143668 | 13591631 |
| 8 | 19588532 | 20790826 |
| 11 | 45034 | 1328434 |
| 11 | 10223062 | 11066107 |
| 12 | 51951428 | 53642447 |
| 13 | 57315001 | 58896270 |
| 14 | 34045334 | 35873987 |
| 15 | 43986135 | 46685544 |
| 16 | 32235312 | 34667199 |
| 17 | 73315353 | 83593846 |
| 18 | 44877843 | 52524300 |
| 19 | 20749258 | 24123054 |
| 20 | 42926137 | 50207630 |
| 21 | 37280554 | 41604383 |
| 22 | 44003464 | 45115762 |
| 23 | 17989759 | 19190868 |
| 23 | 32517742 | 33226380 |
| 24 | 32151959 | 34111200 |
| 25 | 16615782 | 17748902 |
| 26 | 5238353 | 5934769 |
| 26 | 22234700 | 23734968 |
| 27 | 31239942 | 32930888 |
| 28 | 6252554 | 6476001 |
| 28 | 12164625 | 12529021 |
| 29 | 1111747 | 2754119 |

Table S5. Names of gene features and their position within the region of Ssa18. Gene annotation data are based on the newest assembly for Atlantic salmon (Ssal_v3.1).

| **Protein name** | **start (bp)** | **end (bp)** | **LOC** | **Biotype** |
| --- | --- | --- | --- | --- |
| ectonucleoside triphosphate diphosphohydrolase 1 [Salmo salar] | 49505869 | 49546165 | LOC106577463 | protein_coding |
| NA | 49554721 | 49554793 | trnav-uac-69 | tRNA |
| NA | 49554928 | 49555000 | trnav-cac-65 | tRNA |
| NA | 49555744 | 49555816 | trnav-uac-70 | tRNA |
| NA | 49581687 | 49581840 | LOC123728854 | rRNA |
| collagen alpha-1(XXV) chain [Salmo salar] | 49593642 | 50050089 | LOC106577460 | protein_coding |
| ethanolamine-phosphate phospho-lyase [Salmo salar] | 50054440 | 50087446 | etnppl | protein_coding |
| oligosaccharyltransferase complex subunit ostc [Salmo salar] | 50095386 | 50139298 | LOC106577458 | protein_coding |
| 60S ribosomal protein L34 [Salmo salar] | 50145103 | 50150788 | rpl34 | protein_coding |
| sodium/potassium/calcium exchanger 2-like [Salmo salar] | 50199880 | 50383211 | LOC106577456 | protein_coding |
| N-acyl-aromatic-L-amino acid amidohydrolase (carboxylate-forming) B [Salmo salar] | 50535395 | 50538803 | LOC106577455 | protein_coding |
| uncharacterized protein LOC106577454 [Salmo salar] | 50604077 | 50622329 | LOC106577454 | protein_coding |
| NA | 50661835 | 50671185 | LOC106577453 | lncRNA |
| uncharacterized protein LOC100196495 [Salmo salar] (gene_synonym=bola2) | 50738985 | 50741243 | zgc:112271 | protein_coding |
| structure-specific endonuclease subunit SLX1 [Salmo salar] | 50741675 | 50751218 | slx1b | protein_coding |
| uncharacterized protein LOC106577451 [Salmo salar] | 50751206 | 50753756 | LOC106577451 | protein_coding |
| inhibitor of growth protein 2 [Salmo salar] | 50762456 | 50779943 | LOC106577449 | protein_coding |
| leucine-rich repeat and calponin homology domain-containing protein 4-like [Salmo salar] | 50836766 | 50913651 | LOC106577444 | protein_coding |
| NA | 50913660 | 50947627 | LOC106577448 | lncRNA |
| NA | 50952867 | 50958707 | LOC106577447 | lncRNA |
| uncharacterized protein LOC106577443 isoform X1 [Salmo salar] | 51053746 | 51088035 | LOC106577443 | protein_coding |
| LOW QUALITY PROTEIN: NACHT and WD repeat domain-containing protein 2-like [Salmo salar] | 51105966 | 51128419 | LOC106577465 | protein_coding |
| piggyBac transposable element-derived protein 4-like [Salmo salar] | 51487717 | 51488505 | LOC106577389 | protein_coding |
| NA | 52216470 | 52217160 | LOC123728642 | lncRNA |
| NA | 52220196 | 52220886 | LOC123728643 | lncRNA |
| NA | 52224458 | 52225148 | LOC106566019 | lncRNA |
| NA | 52228369 | 52229059 | LOC123728644 | lncRNA |
| NA | 52238651 | 52239341 | LOC123728645 | lncRNA |
| piggyBac transposable element-derived protein 4-like [Salmo salar] | 52255163 | 52255777 | LOC106569786 | protein_coding |
| protocadherin-7-like [Salmo salar] | 52807898 | 53027936 | LOC106577482 | protein_coding |
| NA | 52853107 | 52856516 | LOC123728723 | lncRNA |

Fig. S1. Principal component analyses (PCAs) of 31 outlier regions detected by *lostruct* based on multidimensional scaling (MDS) axis 2. Data points are coloured by continent for North America (red) and Europe (blue). Chromosome numbers and the genomic position of the regions are provided for each plot. The R package *adegenet* (Jombart, 2008) was used to generate PCA data.

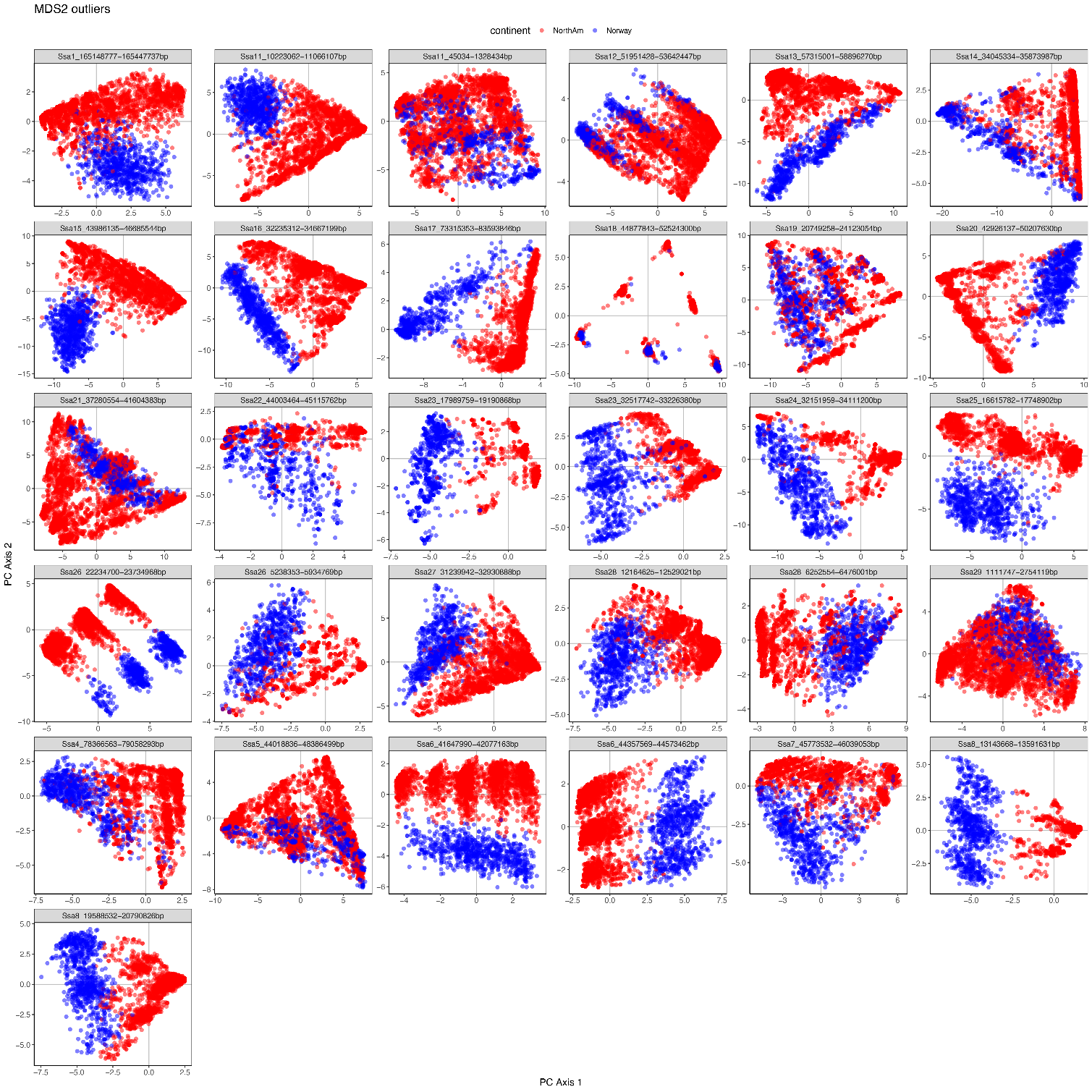

Fig. S2. Pairwise linkage disequilibrium (LD; r^2^) between SNPs on Ssa18 (from 30-60 Mbp) with European values below diagonal and North American above diagonal. High LD is indicated by red and any fixed loci are indicated by white (LD not calculated). The region of high LD identified in the study is indicated by gray bars.

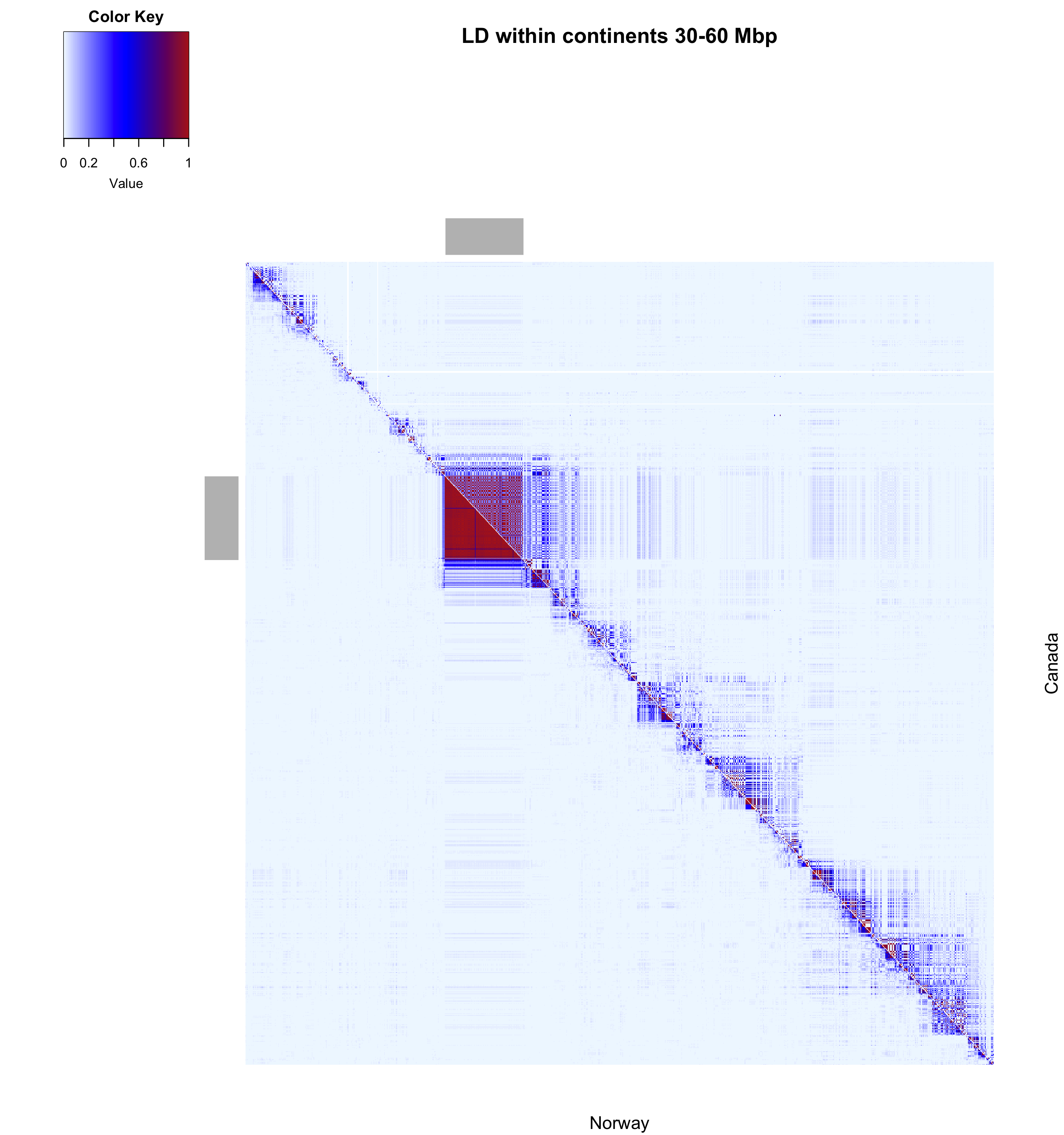

Fig. S3. Principal component analysis (PCA) of Ssa18 region within (A) Europe and (B) North America separately. PCAs were performed using *pcadapt* (Luu et al., 2017).

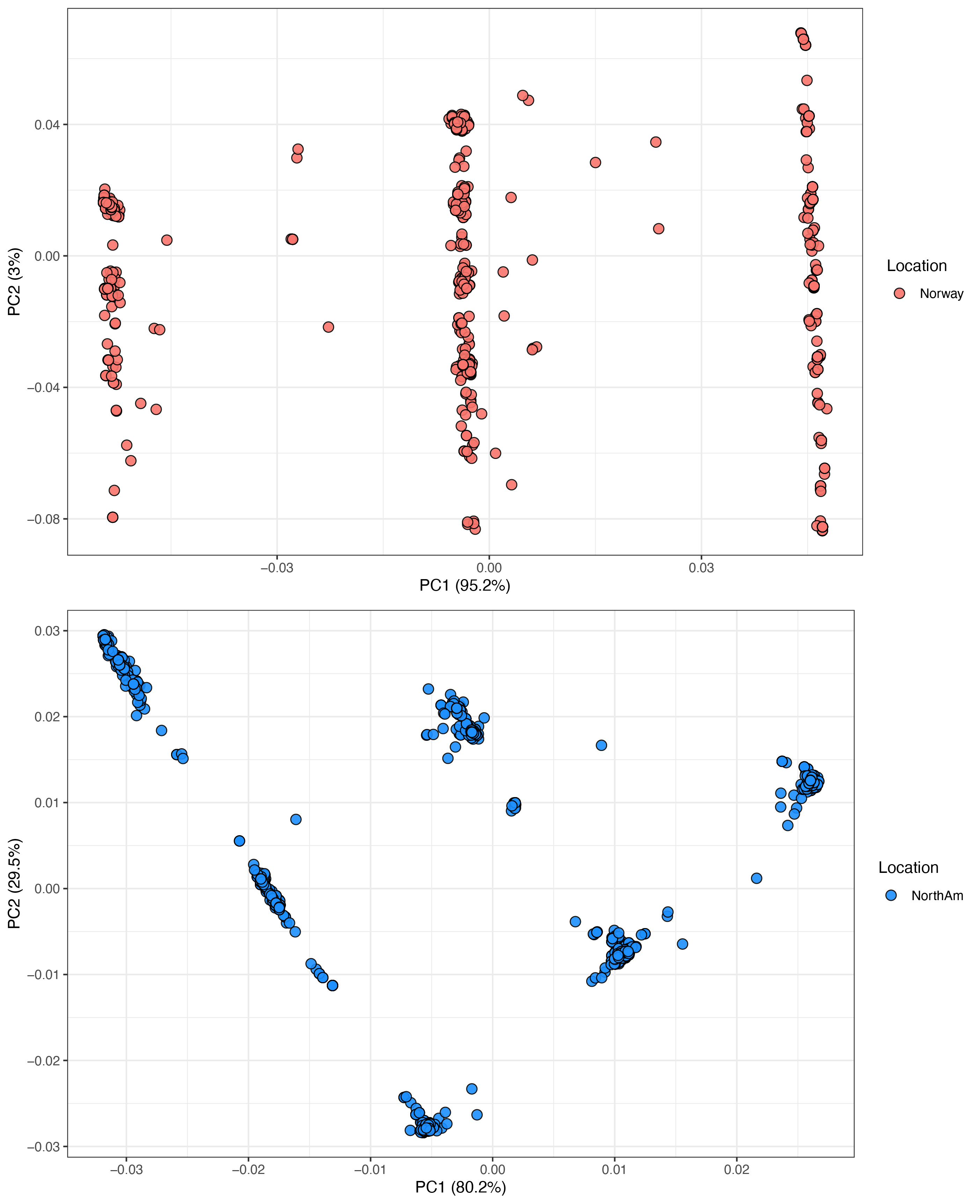

Fig. S4. Map with frequency of the B allele (European origin) in Atlantic salmon (*Salmo salar*) populations from North American sampling sites. We hypothesized that the B allele indicates potential evidence of trans-Atlantic secondary contact (introgression) in the Ssa18 region. Secondary contact was inferred based on clustering patterns in principal component analysis (PCA), where individuals from North America clustered with genotypes that included the European-specific B allele, which included AB, BB, or BC genotypes.

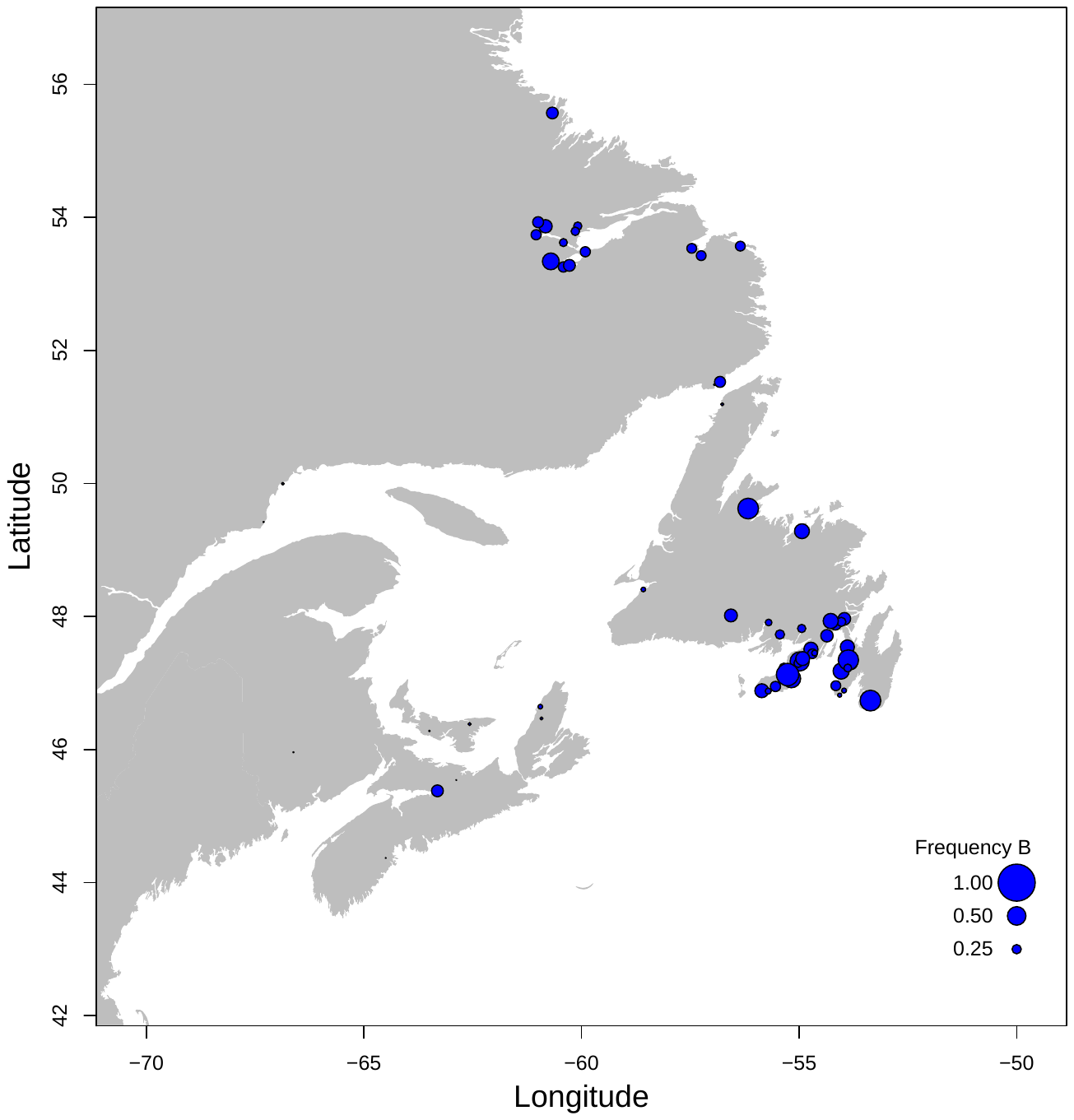

Fig. S5. Relationship between the frequency of the A allele and latitude in (A) North American and (B) European populations of Atlantic salmon (*Salmo salar*). Frequency of A allele was determined through PCA clustering of the haplotype block on Ssa18.

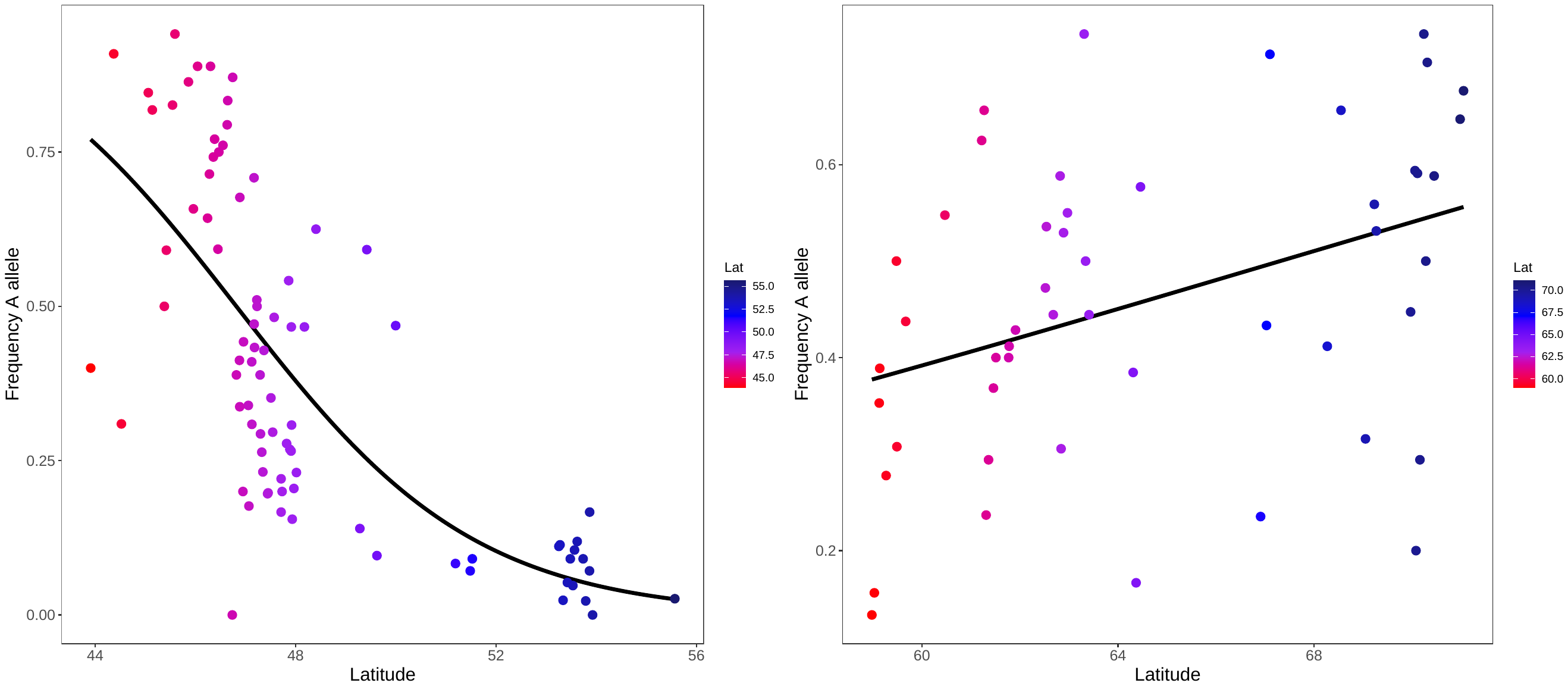

Fig. S6. Relationship between 19 bioclimatic variables (scaled) and haplotype frequency (frequency of the A allele) across North American Atlantic salmon (*Salmo salar*) populations. Smoothed line represents generalized linear relationship generated using ggplot2 R package.

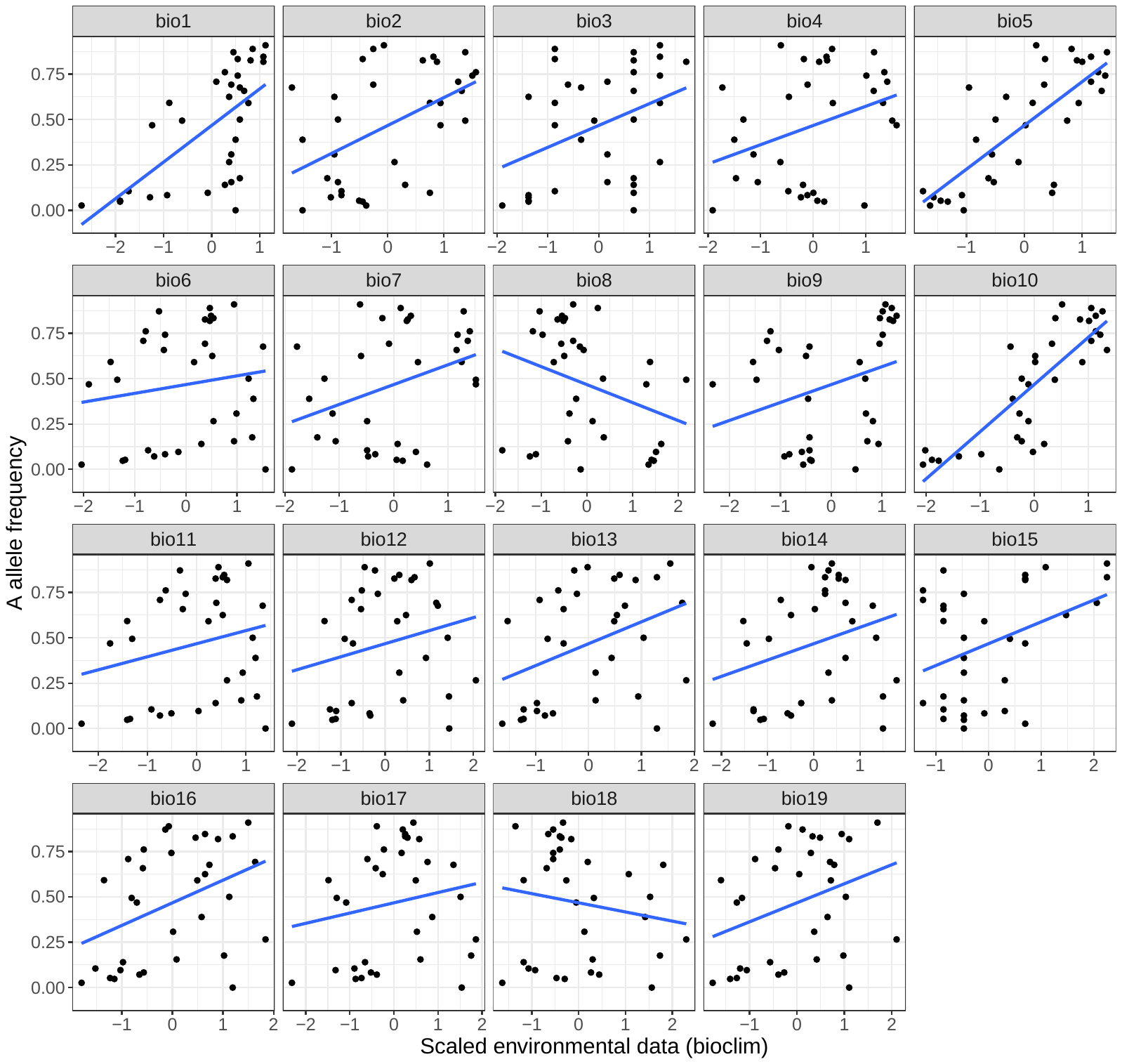

Fig. S7. Relationship between 19 bioclimatic variables (scaled) and haplotype frequency (frequency of the A allele) across European Atlantic salmon (*Salmo salar*) populations. Smoothed line represents generalized linear relationship generated using ggplot2 R package.

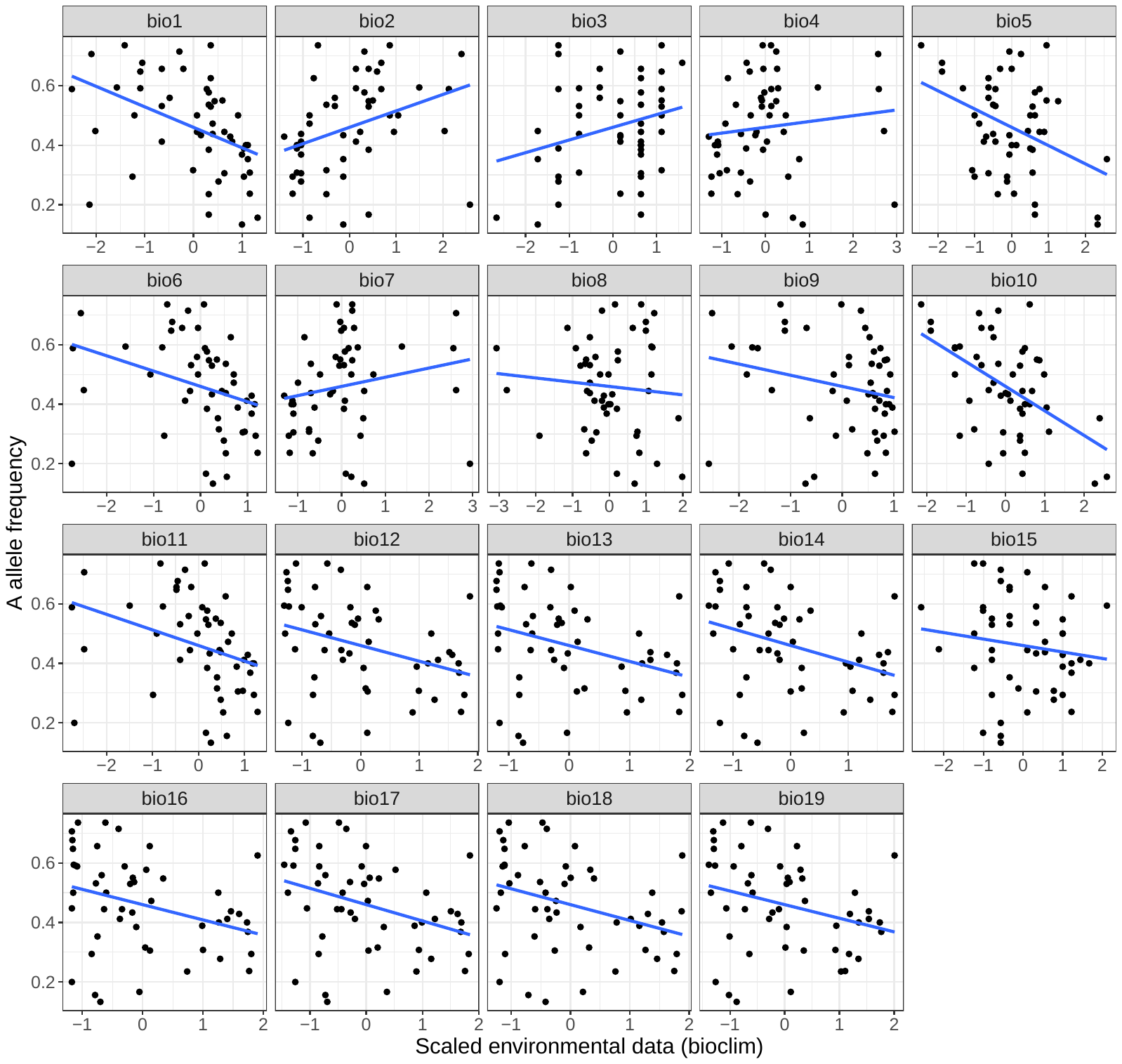
